# Body size and fruit handling ability explain frugivory patterns among Asian mammals

**DOI:** 10.64898/2026.02.24.707853

**Authors:** Bibidishananda Basu, Kim R. McConkey, Rohit Naniwadekar, Sandeep Pulla, Jun Ying Lim, Aparajita Datta

**Affiliations:** Manipal Academy of Higher Education, Manipal, Karnataka; Nature Conservation Foundation, Mysore, Karnataka; BIOTEC, National Science and Technology, Development Agency, Pathom Thani, Thailand; National Centre for Biological Sciences, Bengaluru, Karnataka; National University of Singapore, Singapore

**Keywords:** carnivores, herbivores, frugivores, mammal traits, plant-animal interactions, primates, size-matching hypothesis

## Abstract

**Aim:** Frugivore traits influence the diversity of fruits that frugivores consume. While well studied in birds, trait-based drivers of frugivory remain less understood in mammals despite their considerable morphological, physiological, and ecological diversity. We compiled a comprehensive dataset of mammalian frugivory across Asia, and compared the species diversity and morphological traits of fruits consumed across major mammal guilds. We examine the importance of traits such as body size and fruit handling ability in explaining observed frugivory patterns among Asian mammals.

**Location:** Asia.

**Time period:** 1964–2023.

**Major taxa studied:** ‘Herbivores’ (Perissodactyla, Artiodactyla, Proboscidea), Primates, Carnivora

**Methods:** We compiled frugivory data for three major non-volant, terrestrial Asian mammals from 518 sources, comprising 5276 unique plant–mammal interactions between 142 mammal species and 2,234 plant species. We compared fruit taxonomic composition, diversity and morphological traits (fruit size, colour, type, growth habit, seed number) among three mammal groups. Using Phylogenetic Generalized Least Squares (PGLS) and Ordinary Least Squares (OLS) regression, we tested relationships between mammal body size, fruit handling behaviour and the mean and maximum fruit size consumed by different mammal groups.

**Results:** Focal mammal groups differed in taxonomic diversity of fruits consumed. They also differ in fruit colour, fruit type and the growth habit of fruits consumed. Mammal body size was generally associated with the mean and maximum size of consumed fruits. Fruit handling ability was an important factor for the maximum size of non-fig fruits consumed but not for figs, potentially because figs may be consumed piecemeal.

**Main conclusions:** Our findings highlight the need for a more nuanced understanding of mammalian frugivory patterns due to differences among mammal groups in fruit consumption patterns, range of body sizes and fruit handling abilities. In the context of frugivory interactions, mammals are better considered as different functional groups rather than a single entity.

## 1 | Introduction

Fruits are important, readily accessible energy sources for the animals that feed on them (Fleming & Kress, 2013). Plant-frugivore interactions can either lead to the successful dispersal or destruction of the seed by the animal, both of which are important processes that shape the regeneration, stability, and maintenance of diversity in plant communities across many terrestrial ecosystems (van Leeuwen et al., 2022). The foraging preferences made by frugivores for fruits, and therefore the plant species they consume, are shaped by the traits of both fruits and frugivores (Rojas et al., 2022). Understanding trait associations is important for predicting frugivore diet composition and potential contributions to plant recruitment (McFadden et al., 2022). However, evolution has favoured diffuse interactions between fruits and frugivores (Valenta & Nevo, 2020) because most fruits are consumed by multiple frugivore species with a wide variety of requirements (Fleming & Kress, 2013); therefore fruit traits are under multiple selective pressures (Gautier-Hion et al., 1985).

Trait associations (“trait-matching”) between frugivores and fruit have been studied across a range of characteristics, including fruit colour, pulp reward, seed number and growth habit (Cazetta et al., 2008; Dehling et al., 2016; Flörchinger et al., 2010; Nevo et al., 2018; Rojas et al., 2021; Russo, 2003), the most prominent of which is the relationship between frugivore body size and gape size for birds, with fruit and seed size (Bender et al., 2018; Chen & Moles, 2015; Lim et al., 2020). This relationship emerges partly due to physical constraints on how efficiently a frugivore can swallow or handle increasingly larger fruits and seeds (Dehling et al., 2016; Wheelwright, 1985). However, size alone does not define many observed frugivore-fruit relationships, as some small-sized animals may consume large fruits and seeds (Ong et al., 2022), underscoring the importance of considering suites of traits to fully understand these interactions. Trait-matching studies have also focused disproportionately on birds, where gape size is often identified as a key determinant of seed-swallowing interactions (Zhang et al., 2025), even though mammals are also key frugivores across terrestrial ecosystems (Jordano et al., 2007). Critically, mammals exhibit greater diversity in sensory, physiological, and anatomical traits than birds (Valido et al., 2011), handling fruits in diverse ways beyond solely with the mouthparts, including manipulation using forelimbs or opposable thumbs, meaning that mammal-fruit relationships probably cannot be readily defined by size relationships alone. This complexity raises the fundamental question of whether mammals can be treated as a functionally uniform group: given the trait diversity across mammalian taxa, broad taxonomic groups may differ markedly in their associations with the fruit traits that govern consumption. Hence, understanding how mammals interact with fruit requires a more nuanced trait framework.

To understand the extent to which mammalian groups differ in the species, and trait combinations, of fruits consumed has clear macroecological implications. Fruit species composition and trait distributions vary substantially across vegetation types, as do frugivore assemblages (Fricke et al., 2022); yet most studies (Gautier-Hion et al., 1985; Koike & Masaki, 2019) investigate fruit trait and frugivore assemblages in single ecosystems. Community-level studies suggest that sympatric mammal taxa sometimes consume similar fruits (Koike & Masaki, 2019) and at other times their preferences diverge (Forget et al., 2007; Gautier-Hion et al., 1985), but whether this variation is predictable beyond single ecosystems and vegetation gradients is unknown. Therefore, characterizing broad-scale patterns may help to understand the general patterns and the extent of variation (Lim et al., 2020; Dracxler & Kissling, 2022; Simmons et al., 2018). A continental-scale synthesis would allow integration across multiple local studies, capturing spatial variation in fruit–frugivore interactions across vegetation types and biogeographic contexts (Fricke et al., 2022), thereby providing a more comprehensive understanding of mammal–fruit interactions.

The Asian continent harbours nearly a quarter of the world’s terrestrial mammal species (∼1,000; Corlett, 1998), including more than 80 large herbivores (Sridhara et al., 2016), over 130 fruit-eating primates (McConkey, 2018), and 85 carnivores (Fernández-Sepúlveda & Martín, 2022), distributed across a wide range of vegetation types, from tropical rainforests to dry deciduous forests, grasslands, and montane systems. This diversity of frugivore assemblages makes Asia ideally suited to examine patterns of fruit consumption and trait matching across major frugivorous mammal groups. In this paper, we explore the taxonomic overlap in fruit consumption and the variation in fruit traits consumed among three major frugivorous mammal groups in Asia: carnivores, primates, and herbivores (Corlett, 1998). We address the following questions: (1) To what extent do these groups overlap in the taxonomic diversity of fruits consumed? (2) How do fruit morphological traits vary among the three mammal groups that consume them? (3) How are morphological traits of mammalian frugivores, specifically body size, and fruit handling behaviour, associated with the fruit size and they consume?

## 2 | MATERIALS & METHODS

### 2.1 | Interaction data

Our study focuses on three major terrestrial non-volant mammal frugivore groups: carnivores (order Carnivora), primates (order Primates) and herbivores (orders Artiodactyla, Perissodactyla and Proboscidea).

We compiled data on ripe fruit consumption for the focal mammal groups using Google Scholar. We used multiple keyword combinations for searches, including the scientific and common names of each mammal along with terms ‘diet ecology,’ ‘feeding ecology,’ ‘seed dispersal,’ and ‘frugivory.’ For each combination of terms, we obtained studies up to the tenth results page. Each paper was checked manually for information about ripe fruit consumption: i.e., mentioned explicitly as ripe fruit handling, or intact seed deposition. From each valid study, we obtained a list of ripe fruit species consumed by the studied mammal species.

In total, we collected fruit consumption data for 32 carnivore species (24 genera), 38 herbivore species (24 genera), and 72 primate species (16 genera). We reviewed 518 papers from 19 Asian countries (Fig. 1). We digitized 8,732 mammal-frugivore interaction records, encompassing 2,234 plant species (639 genera and 120 families) (Electronic supplementary material 1). We did not include any unidentified plant species in the analysis. A list of data sources used in the study are provided in the data sources section. Fruit consumption data for domesticated animals (11 species) were included for overlap of taxonomic diversity and differences in morphological traits among mammal groups but were excluded from our frugivore body size and fruit size analyses.

**Figure 1.**
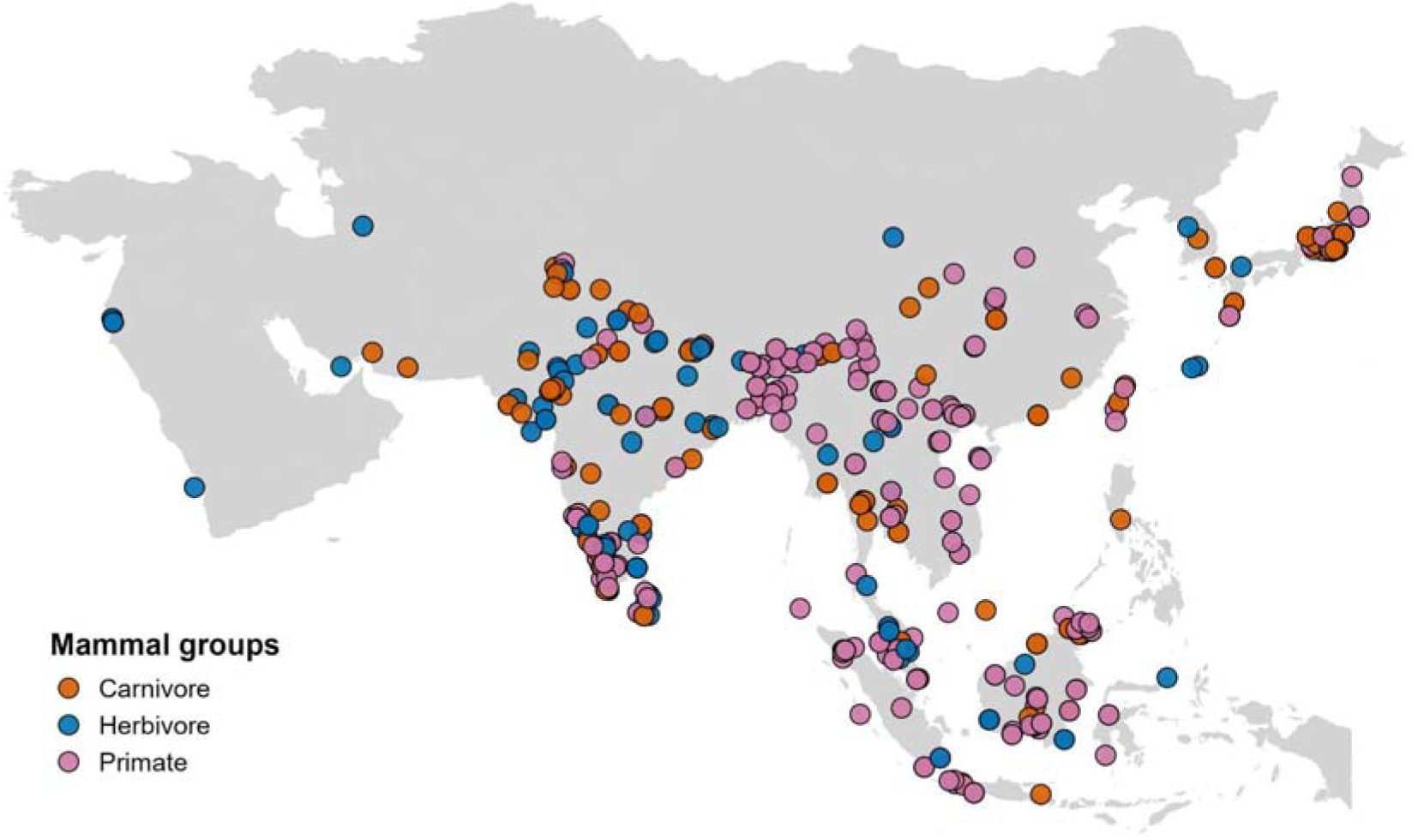
Map of Asia showing the study locations (n = 518 studies) of mammal species.

### 2.2 | Plant Traits

We compiled fruit trait data from 195 sources (Electronic supplementary material 2), including floras, books, and research articles. The dataset includes six morphological traits: growth habit (tree, shrub, herb, climber), fruit type (dry, fleshy), number of seeds per fruit, fruit length and diameter, and ripe fruit colour. For growth habit, the only herbaceous plants were members of the Musaceae.

We categorized fruits into figs and non-figs for traits such as fruit length, diameter and colour; for all other traits, figs were not treated separately. For the frugivore body size and fruit size relationship analyses, we treated them separately. Mammals, and other frugivores, may interact differently with fig and non-fig fruits because these fruit types differ markedly in morphology and nutritional characteristics (O’Brien et al., 1998; Shanahan et al., 2001). Figs typically contain numerous minute seeds, are soft, fleshy with no outer covering/skin that needs to be removed or handled, and can be consumed piecemeal more easily than many other fruits. Figs also frequently exhibit asynchronous fruiting, have large crop sizes, and relatively short fruit-ripening durations (Lambert & Marshall, 1991). These traits make figs important to frugivores in many habitats. Collectively, morphological traits with other characteristics provide a strong rationale for treating figs separately from other fruit groups in our analyses.

### 2.3 | Mammal Traits

We compiled trait data for 131 mammal species focusing on characteristics relevant to frugivory: body size (male and female weight ranges) (Electronic Supplementary Material 3). In addition, the fruit handling ability of each species was classified into three broad categories (“mouthparts only”, “forelimbs”, “opposable thumbs”) informed by observations of fruit handling behaviour (see Electronic Supplementary Material 4). We identified many carnivores and herbivores (e.g., cervids) as handling fruits exclusively with their mouthparts (Iwaniuk & Whishaw, 2000). By contrast, we identified some carnivores, including viverrids, herpestids, and ursids, that can additionally use their forelimbs to grasp and manipulate fruits, with greater motor control and flexibility in their forelimbs allowing more effective manipulation during feeding. Lastly, we included all of the region’s primates in the final category as they all possess opposable thumbs, which confers even more dexterity for fruit consumption (Fragaszy & Crast, 2016). We hypothesise that these categories of fruit handling ability will influence the range of fruits that mammals can effectively manipulate and consume.

### 2.4 | Statistical Analyses

We verified and updated all plant scientific names using Plants of the World Online (https://powo.science.kew.org/). All analyses were conducted in R version 4.2.2. We quantified taxonomic overlap using percentage overlap with unique mammal species and plant species combination. Differences in the genera and families consumed by the three mammal groups were tested using PERMANOVA (adonis function, *vegan* package) (Dixon, 2003) based on Bray–Curtis dissimilarities calculated from a binary presence–absence matrix, with 999 permutations.

To examine variation in continuous fruit traits (fruit length, fruit diameter, and seed number per fruit), we calculated mean trait values for each mammal species and reported group-level means with bootstrapped 95% confidence intervals. Differences among mammal groups were tested using Kruskal-Wallis test and pairwise Wilcoxon rank-sum tests, with *p*-values adjusted for multiple comparisons using the Benjamini-Hochberg procedure.

Fruit colours were grouped into three categories based on contrast against a green leaf background (Nevo et al., 2018): bright (red, orange, pink), dark (black, blue, purple, violet), and dull (brown, green, yellow, grey, white). For categorical fruit traits (colour, growth habit, and fruit type), we tested associations with mammal groups using Chi-square tests for independence and we also measured differences between mammal groups with the same. For activity pattern and fruit colour consumed by each mammal species, we determined the significance of these associations through Chi-square tests for independence. Vegetation types were classified into two broad categories, Temperate & Subtropical and Tropical, using GPS coordinates from each study and biome information (Dinerstein et al., 2017). Ten biomes with several ecoregions were consolidated into these two categories (Table S1). For each site, the GPS locations and study description were reviewed to determine and verify the accuracy of vegetation type assignment. Around 3 sites out of 371 were re-classified based on specific site-level vegetation and climatic regime information.

We used the iNEXT R package (Hsieh et al., 2016) to generate species accumulation curves based on Hill diversity indices and to assess sampling coverage for each mammal group. We used unique mammal species and plant species combinations (5276 interactions) to ensure we did not over-represent interactions from single sites with multiple publications.

To examine the relationship between fruit diameter and mammal body mass across handling types, we performed OLS and PGLS regression analysis. Mean and maximum fruit diameters were evaluated as separate response variables. To assess the relative importance of fruit handling ability, log-transformed mammal body mass, and their interaction, we compared four candidate models for each response variable: (1) a null intercept model (y∼1), (2) a body size model y ∼ log (Body size), (3) a fruit handling model (y∼fruit handle), and (4) an interaction model (y∼ fruit handle * log (Body size)). Candidate models were evaluated and compared using the Akaike Information Criterion (AIC). We used this analysis for mammal species with ≥10 interaction records..

We performed PGLS to account for the shared evolutionary history among species, with log-transformed mammal body size as the predictor variable and maximum fruit diameter as the response variable. We used 1000 phylogenetic trees based on the mammalian supertree provided by Faurby et al., (2018). Phylogenetic pruning was carried out using the drop.tip() function from the ape package in R, applied across a set of phylogenetic trees stored as a multiPhylo object. PGLS models using maximum likelihood using the phylolm function. To get different parameters for different models in PGLS, we used median value across 1000 phylogenetic trees. This analysis was also interpreted in the context of fruit handling ability among mammal groups with the same predictors and response variables.

All analyses were conducted in R version 4.2.2. All the data and code are submitted with the manuscript.

## 3 | RESULTS

### 3.1 | Taxonomic richness and overlap among mammal groups

Primates (72 species) consumed 1786 fruit species (sample coverage: 77.5%) belonging to 631 genera (96%), and 134 families (99%). Herbivores (40 species) consumed 349 species (64.92%) belonging to 185 genera (85.2%), and 64 families (97%). Carnivores (30 species) consumed 717 species (70.83%) belonging to 325 genera (92%), and 103 families (98%). The species accumulation curves indicate (Fig. S1) that herbivore dietary data are less complete than those for primates and carnivores. Across vegetation types, taxonomic richness of consumed fruit species shows similar trends (Fig. S2 & Table S2).

Across Asia and its various vegetation types, the composition of fig, non-fig fruit genera and species, and families (Table 1 & Table S3-S4) consumed by the three mammal groups differs significantly (see Table 1 for PERMANOVA results). Primates consume more unique fruit species across Asia, regardless of vegetation type, fig and non-fig fruits. Fruits eaten by herbivores are largely shared with the other groups. For non-fig fruit genera and families, there is greater overlap between the fruits consumed by carnivores and primates (Table 1).

**Table 1.** Fruit consumption by carnivores, herbivores, and primates across Asia. Values represent the percentage of plant species, genera and families consumed exclusively by each group or shared between two or all three groups, analysed separately for non-fig genera, and for fig species. The final column presents *p*-values from PERMANOVA tests for mammal group-level differences in consumption.

|  | **Sample size** | **Uniquely eaten (%)** |  |  | **Shared (%)** |  |  |  | **PERMANOVA *p*-value** |
|---|---|---|---|---|---|---|---|---|---|
|  |  | **Herbivore** | **Carnivore** | **Primate** | **Carnivore & Primate** | **Carnivore & Herbivore** | **Herbivore & Primate** | **All Groups** |  |
| **Fig species** | **114** | **4** | **11** | **46** | **20** | **2** | **5** | **13** | **0.001\*** |
| **Non-fig species** | **2120** | **5** | **14** | **58** | **12** | **1** | **5** | **5** | **0.001\*** |
| **Non-fig genus** | **721** | **2** | **9** | **44** | **21** | **6** | **8** | **15** | **0.001\*** |
| **Non-fig family** | **148** | **2** | **5** | **23** | **26** | **1** | **4** | **38** | **0.001\*** |

### 3.2 | Functional fruit trait variation among terrestrial mammal groups

Herbivores eat larger non-fig fruits, although mean fruit diameter did not differ significantly among mammal groups (Fig. 2, Table S5). The size of non-fig fruits eaten by carnivores is more similar to those consumed by primates than by herbivores (Fig. 2a & 2b). For figs, the mean fruit size consumed was broadly similar across groups, except that herbivores consumed significantly larger fruit diameters than the other groups at the Asia level (Fig. 2b).

**Figure 2.**
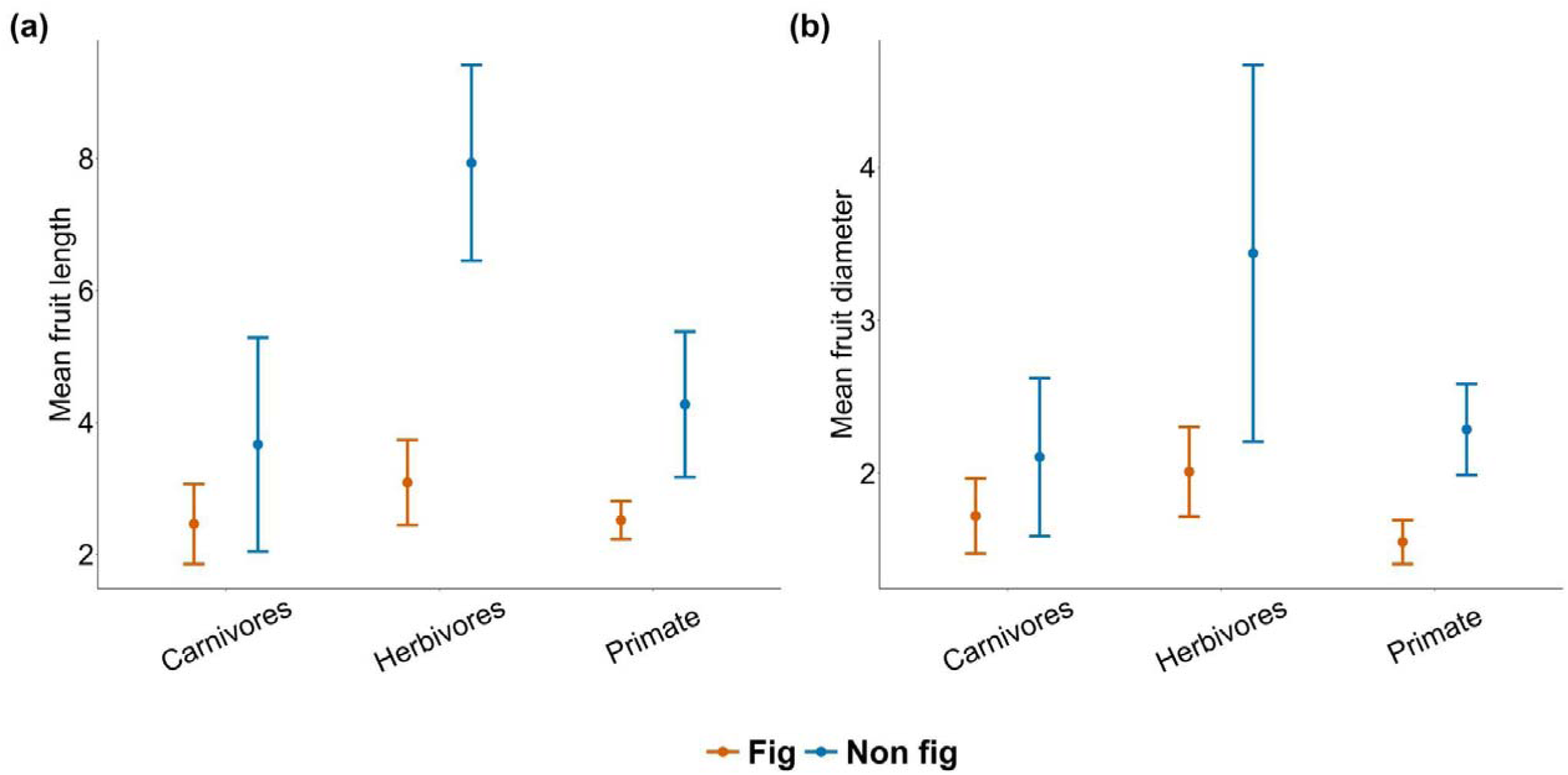
Differences in mean fruit length and mean fruit diameter (fig and non-fig species) consumed by focal mammal groups across Asia (a & b).; for a, fruit length for fig (Carnivore: 30 interactions, Herbivore: 17, Primate: 76), for non-fig (Carnivore: 698, Herbivore: 535, Primate: 2186); For b, fruit diameter of fig (Carnivore: 93, Herbivore: 55, Primate: 424), for non-fig (Carnivore: 1027, Herbivore: 615, Primate: 2848). Error bars represent 95% confidence intervals.

Colour of figs (χ² = 11.06, df = 4, *p* = 0.02; Fig. 3a) and non-fig fruits (χ² = 120.52, df = 4, *p* < 0.01) consumed also differed significantly among mammal groups. For non-fig fruits, carnivores consumed across all colour categories in similar proportion, whereas herbivores and primates mainly consumed dull-coloured fruits (Fig. 3b.). Pairwise comparison reveals that, for figs there is a difference between fruit colors consumed by herbivores and primates and the three groups differ significantly with non-fig colour they consume (Table S6).

**Figure 3.**
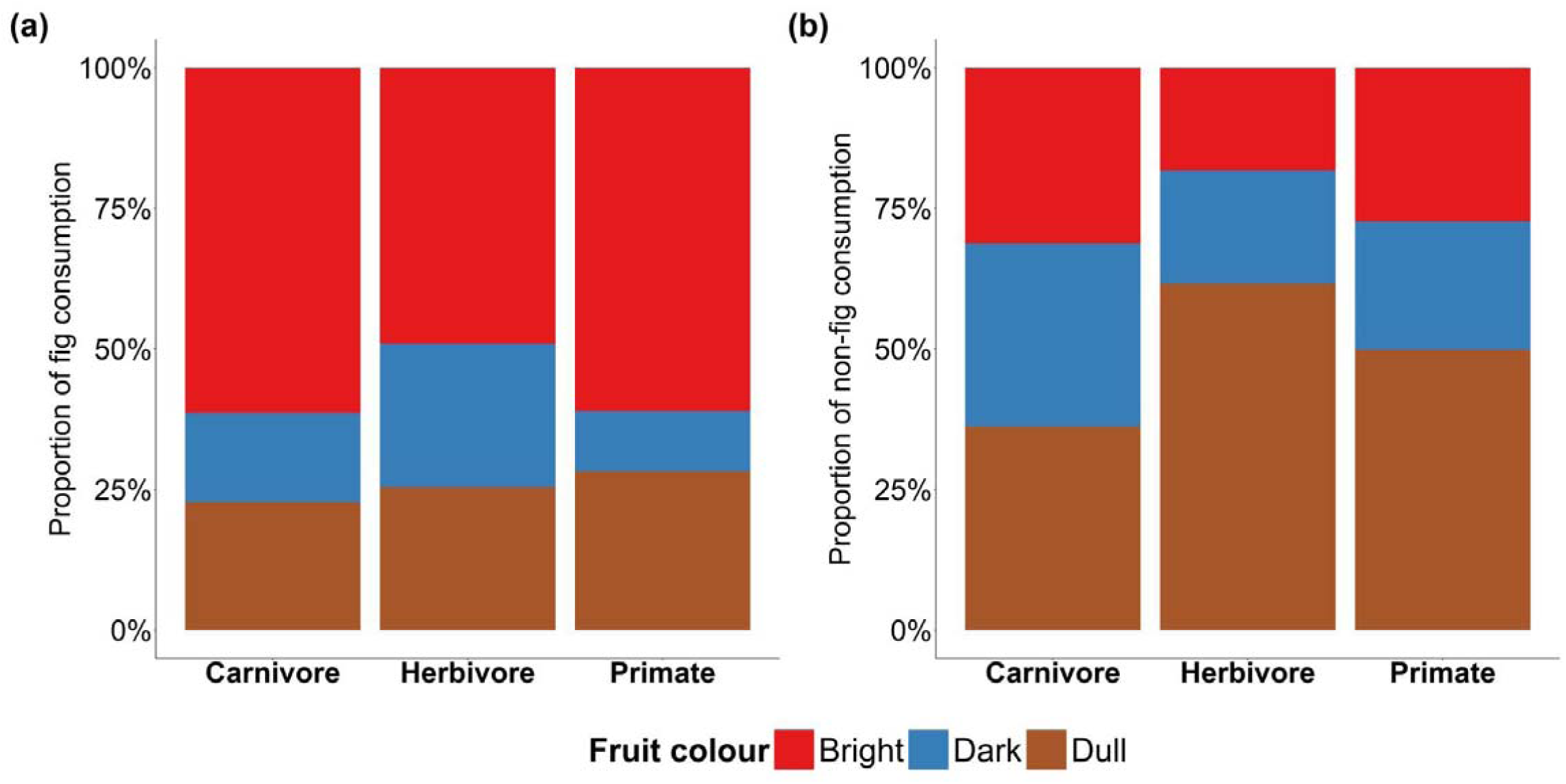
Fruit colours of fig and non-fig fruits consumed by different mammal groups across Asia. (a) colour pattern of fig fruits (Carnivores: 101 interactions, Herbivores: 55, Primates: 429) & (b) non-fig fruits (Carnivores: 1099, Herbivores: 641, Primates: 3028) across Asia.

The growth habits of fruiting plants consumed by the focal mammal groups differ significantly across Asia (χ² = 615.54, df = 4, *p* < 0.01). Herbivores consume fruits from more trees than do carnivores and primates across Asia (Fig. 4a & 4b). Carnivores feed on more climbers and shrubs than the other groups. Pairwise comparison reveals that all the mammal groups differ in the consumption of fruits based on growth habit (Table S6).

**Figure 4.**
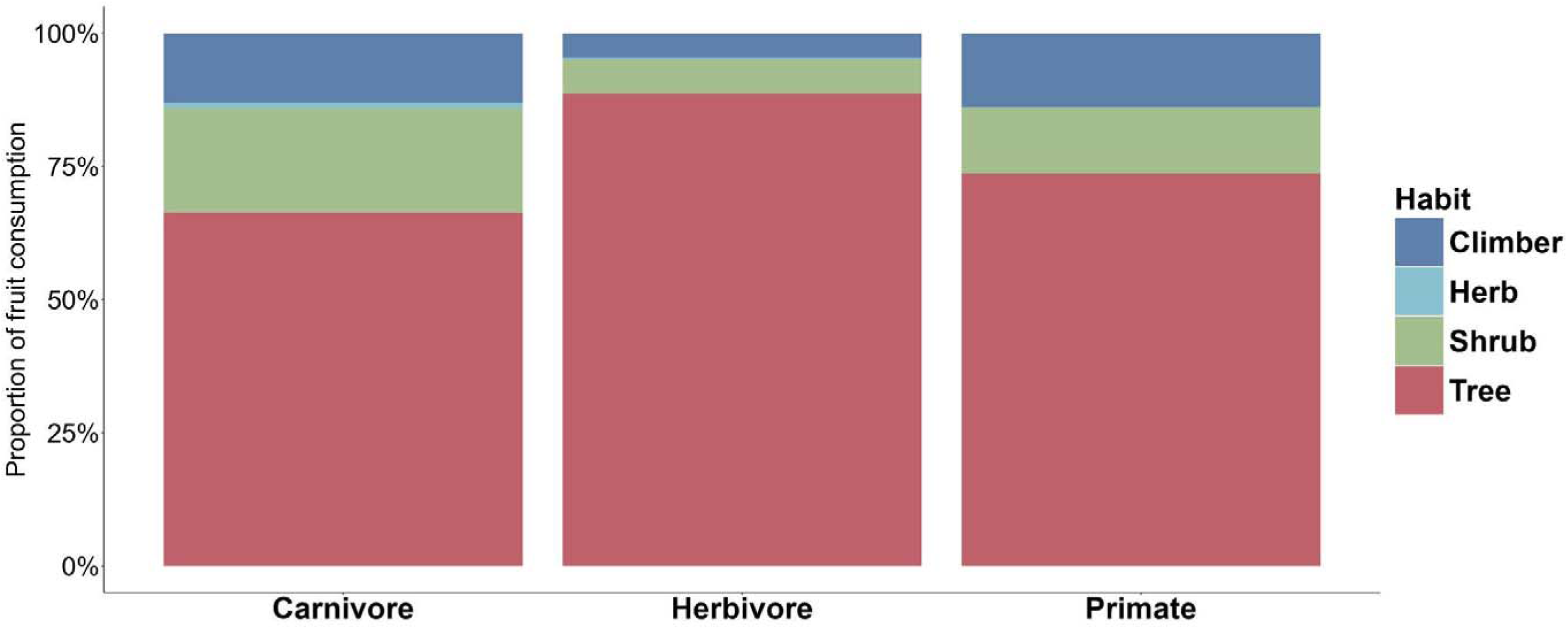
Growth habit of the fruits consumed by different mammal groups across Asia. (Carnivore: 1204 interactions, Herbivore: 699, Primate: 3465).

The fruit type of plants consumed by the three mammal groups differ significantly across Asia (χ² = 25.88, df = 2, *p* < 0.01). Carnivores feed on more fleshy fruits than the other groups (Fig S3). Pairwise comparisons reveal that carnivores differ significantly from other groups in fruit types they are consuming (Table S6).

Mammal groups did not differ significantly in number of seeds per fruit for the fruits consumed by them (Table S5), Herbivores consumed fruits with higher mean number of seeds per fruit than other groups (Fig. S4).

Across all vegetation types, the fruit traits show similar patterns (Table S5 & S6, Figure S5-S9).

### 3.3 | Effect of body size and handling ability on patterns of fruit size consumption

For mean and maximum fig size, the body size and null models performed best (ΔAIC = 0), respectively. For mean and maximum non-fig size, the body size and interaction models performed best (ΔAIC = 0). For both OLS (Fig. 5; Table 2) and PGLS (Fig. S10; Table 2) model performances are similar (Table S7).

**Figure 5.**
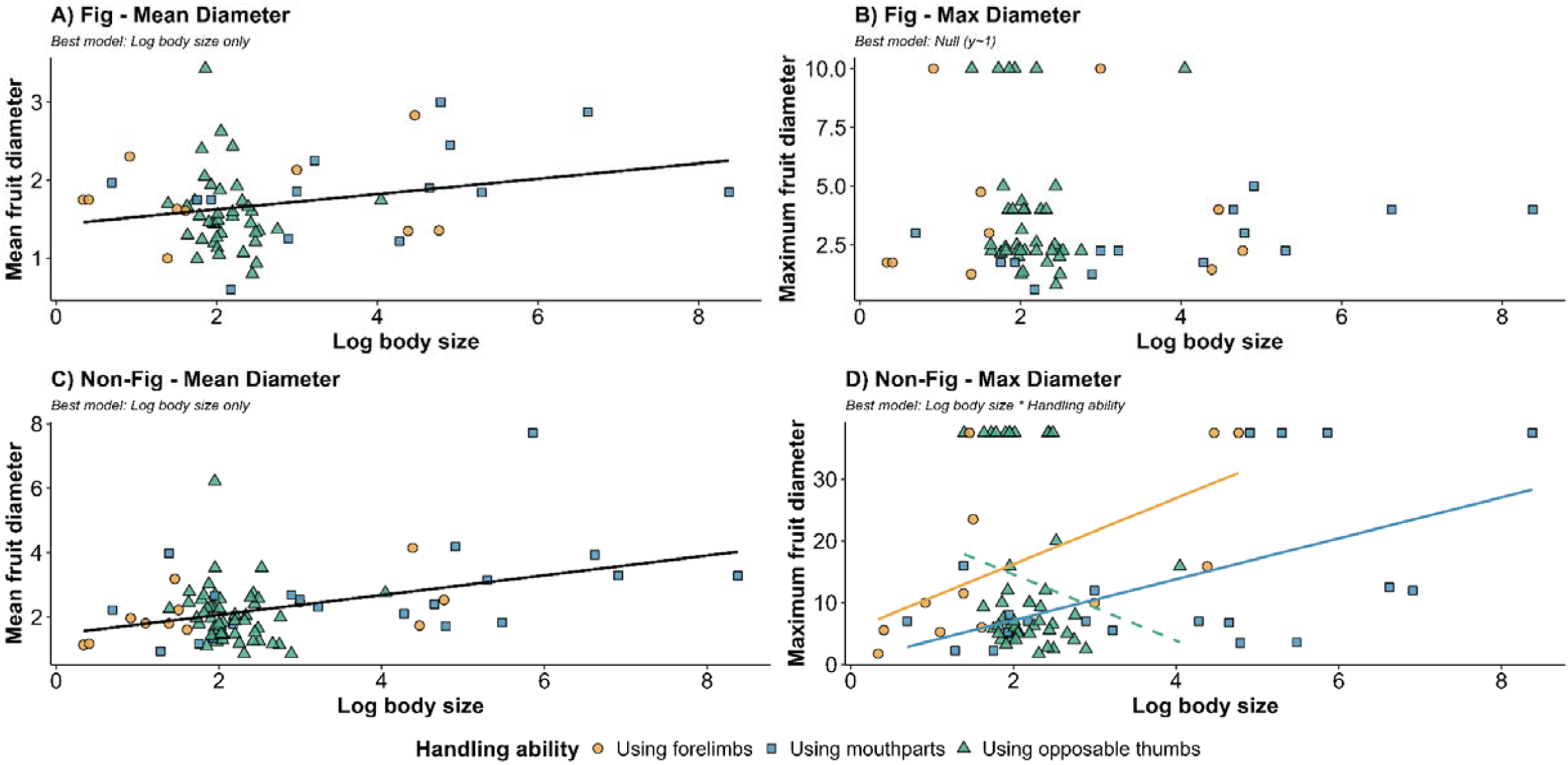
Relationships between mammal body size, fruit handling ability, and consumed fruit diameter. Panels display mean **(A)** and maximum **(B)** fruit diameter for fig species (n = 63), and mean **(C)** and maximum **(D)** diameter for non-fig species (n = 82) plotted against log body size (for species with ≥ 10 interaction records). Points represent mammal species categorized by fruit handling ability (forelimbs, only mouthparts, opposable thumbs). Lines depict best-fit OLS models selected via AIC, with solid lines indicating significant relationships (p < 0.05) and dashed lines denoting non-significant trends (p > 0.05).

**Table 2.**
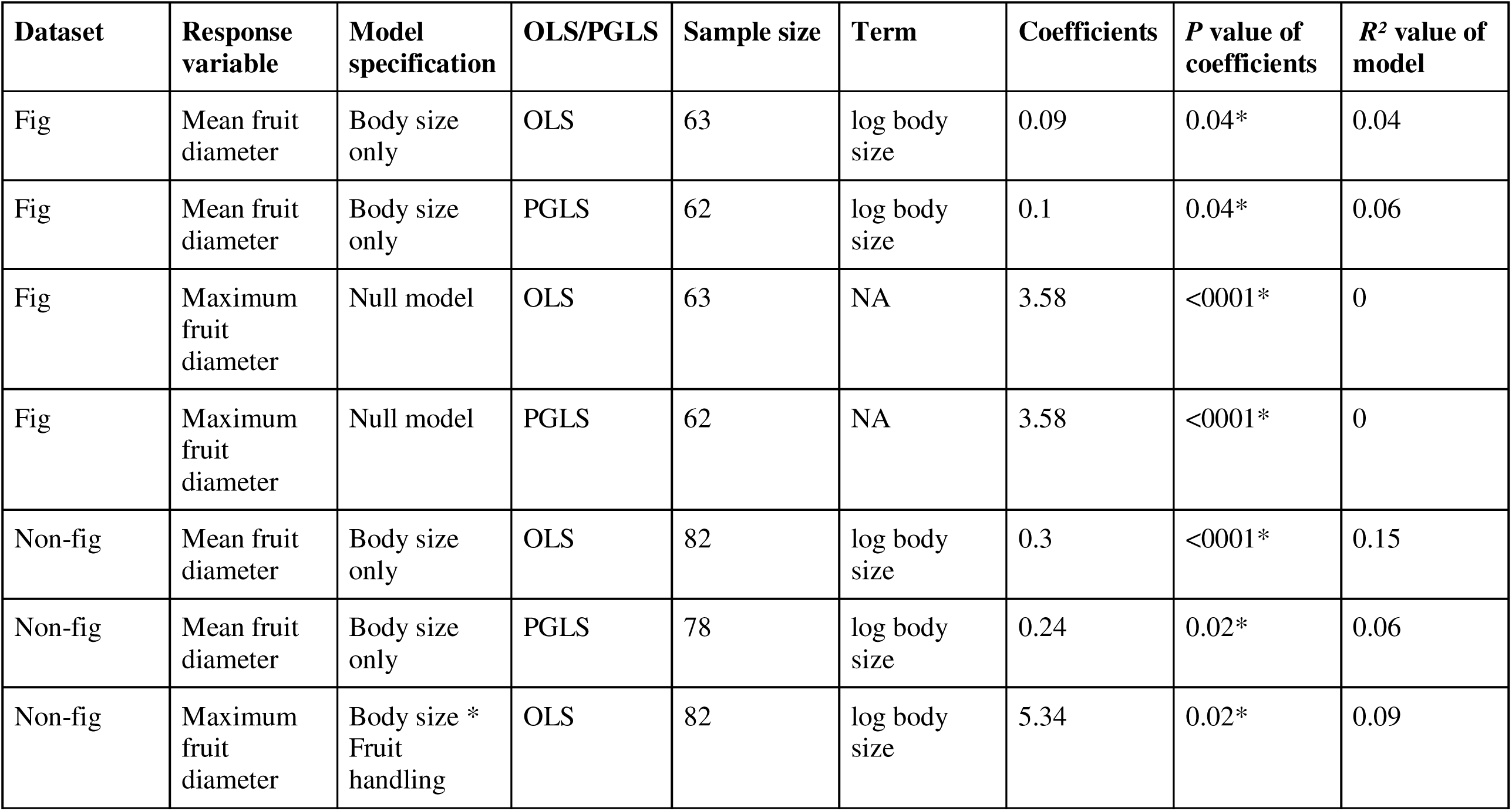
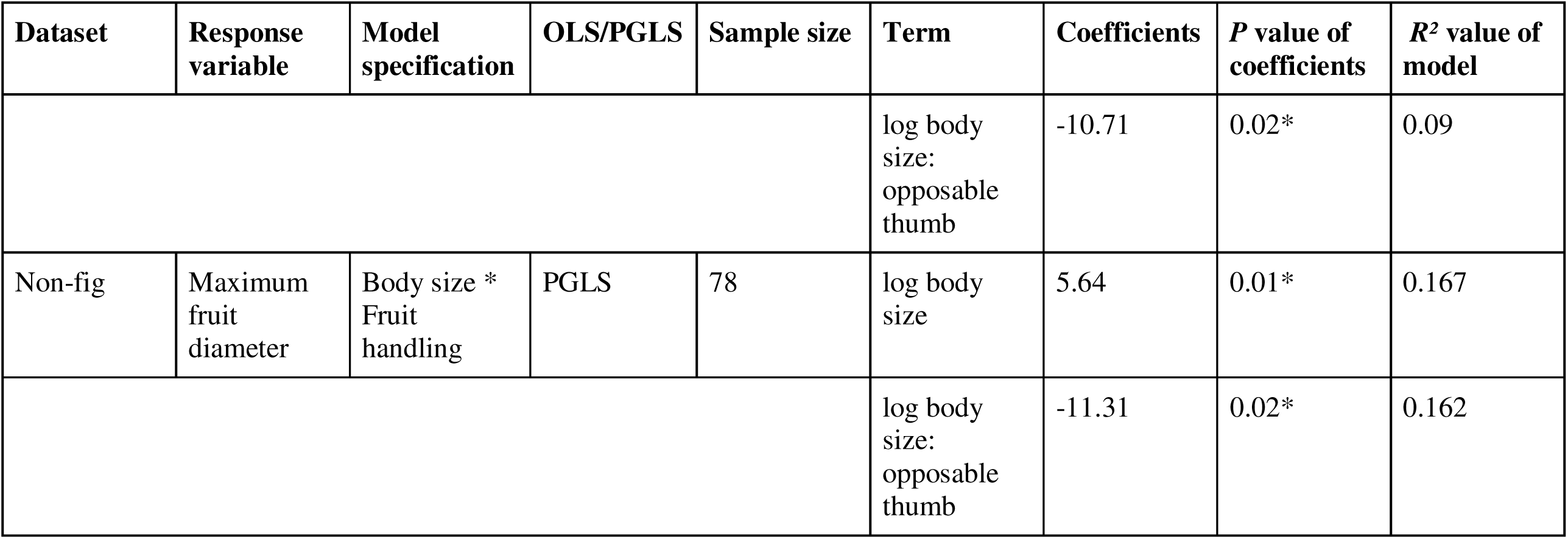
Best supported regression models (ΔAIC = 0) for mean and maximum fruit size of fig and non-fig fruits using OLS and PGLS. For PGLS models, *p*-values and R² values are reported as the median across 1,000 phylogenetic trees. Analyses were restricted to frugivore species with ≥ 10 interaction records. Only coefficient values for significant variables are shown.

| **Dataset** | **Response variable** | **Model specification** | **OLS/PGLS** | **Sample size** | **Term** | **Coefficients** | ***P* value of coefficients** | ***R*<sup>2</sup> value of model** |
|---|---|---|---|---|---|---|---|---|
| Fig | Mean fruit diameter | Body size only | OLS | 63 | log body size | 0.09 | 0.04* | 0.04 |
| Fig | Mean fruit diameter | Body size only | PGLS | 62 | log body size | 0.1 | 0.04* | 0.06 |
| Fig | Maximum fruit diameter | Null model | OLS | 63 | NA | 3.58 | <0001* | 0 |
| Fig | Maximum fruit diameter | Null model | PGLS | 62 | NA | 3.58 | <0001* | 0 |
| Non-fig | Mean fruit diameter | Body size only | OLS | 82 | log body size | 0.3 | <0001* | 0.15 |
| Non-fig | Mean fruit diameter | Body size only | PGLS | 78 | log body size | 0.24 | 0.02* | 0.06 |
| Non-fig | Maximum fruit diameter | Body size * Fruit handling | OLS | 82 | log body size | 5.34 | 0.02* | 0.09 |
| **Dataset** | **Response variable** | **Model specification** | **OLS/PGLS** | **Sample size** | **Term** | **Coefficients** | ***P* value of coefficients** | ***R*<sup>2</sup> value of model** |
|  |  |  |  |  | log body size: opposable thumb | -10.71 | 0.02* | 0.09 |
| Non-fig | Maximum fruit diameter | Body size * Fruit handling | PGLS | 78 | log body size | 5.64 | 0.01* | 0.167 |
|  |  |  |  |  | log body size: opposable thumb | -11.31 | 0.02* | 0.162 |

For mean fruit size, log body size showed a weak positive relationship with fruit size for both figs (OLS: *R²* = 0.06, *p* = 0.04; PGLS: *R²* = 0.06, *p* = 0.04) and non-figs (OLS: *R²* = 0.157, *p* < 0.0001; PGLS: *R²* = 0.06, *p* = 0.01) for body size models. For maximum fruit size, in the interaction model, log body size showed a weak positive relationship with maximum fruit size for non-fig fruits (OLS: *R²* = 0.09, *p* = 0.02; PGLS: *R²* = 0.16, *p* = 0.01) (Table 2). In contrast, there was no significant relationship between mammal body size and maximum fig fruit size (Fig 5, S10).

Although fruit handling ability did not show a significant relationship with fruit size when considered alone, the interaction model indicated positive associations between body size and maximum non-fig fruit size for mammals that handle fruits using only their mouthparts and those that use their forelimbs. The positive relationship between body size and maximum non-fig fruit size was significant across both OLS and PGLS for mammals that handle fruits using only their mouthparts (Table S10).

## 4 | Discussion

Our findings suggest that mammalian groups differ in the diversity of fruits they consume and in some morphological traits of those fruits. We found that body size is an important predictor of the mean fruit size consumed, whereas both body size and fruit-handling behaviour are important predictors of the maximum fruit size consumed. These findings suggest that mammals exhibit substantial diversity in body size, dietary preferences, and fruit-handling abilities, potentially more than observed in birds. Consequently, mammals should not be considered as a single, uniform group when examining frugivory interactions.

### 4.1 | Mammal frugivores vary in taxonomic composition of fruits consumed

Primates are major frugivores in our dataset, consuming many fruit taxa not consumed by the other mammals, reflecting the mainly frugivorous diets of many Asian primates, such as macaques and gibbons (McConkey, 2018). In contrast, carnivores do not rely primarily on fruit; their consumption is often seasonal, with shifts between animal prey and plant resources (Ben-David et al., 1997; Koike & Masaki, 2019) and, consequently, our finding of high diversity of fruits in the diet of some carnivores is an important finding. Carnivores such as civets, bears, and martens, along with primates, can feed on fruits both in the canopy and on the ground. This shared access to multiple foraging strata, and their tendency to consume similar fruits across regions (McConkey & Brockelman, 2011; Séguigne et al., 2022), may explain the higher percentage of fruits shared between carnivores and primates. Herbivore-consumed fruits are mostly shared by other two groups and their low fruit consumption patterns might be an artifact of low sample size compared to others. The differences in the composition of fruits consumed are possibly a reflection of differences in fruit morphological traits consumed by each mammal group. This indicates that there might be complementarity in frugivory between these focal mammal groups.

### 4.2 | Fruits consumed by different mammal frugivores vary in their functional traits

The focal mammal groups differed from each other in multiple traits indicating that mammal groups are functionally different from each other. Most of the mammals covered in this study are important seed dispersers (McConkey, 2018; Nakashima & Do Linh San, 2022; Sridhara et al., 2016). Given that mammal groups differ in both the composition of fruit species they consume and the morphological traits of those fruits, these differences may contribute to the complementarity of seed dispersal functions within plant communities across Asia, as different mammal functional groups may disperse fruits with distinct morphological characteristics. This highlights the importance of these mammals for the regeneration and persistence of mammal-dispersed plant species, particularly given that many of these mammal species are threatened.

Herbivores and primates consume dull-coloured fruits (green, brown, yellow, and grey) more frequently than carnivores. This pattern is well documented for herbivores (Sridhara et al., 2016), as most large herbivores are dichromatic and lack the ability to discriminate between orange and red (Ahnelt & Kolb, 2000; Peichl, 2005), but was unexpected for primates, who are generally known to favour fruits which contrast more from the leaf background (Nevo et al., 2018). However most existing studies have focused on seed-dispersing primates. In contrast, our dataset includes substantial records for langurs and colobine monkeys, which are usually known as seed predators and mainly as folivores (Chen et al., 2023). Seed predators do not rely on cues such as fruit colour or the contrast between fruit colour and the leafy background (Gautier-Hion et al., 1985), features that often function to attract seed dispersers. This difference in foraging behaviour and consumption pattern may explain the observed pattern. Interestingly, carnivores consume bright-and dark-coloured fruits more than other groups. Bright and dark-coloured fruits are typically also bird-dispersed and are often rich in proteins and lipids (Lei et al., 2021). Although carnivores primarily obtain protein and lipids from animal prey, we hypothesize that fruits provide an energetically inexpensive alternative when abundant, may be seasonally important, and may require less foraging effort than hunting for animal prey, leading carnivores to consume more bird fruits than other mammal groups.

Across Asia and vegetation types, mammals generally consumed figs of similar sizes. In our dataset, 48% of fig species are brightly coloured, and all mammal groups consumed more bright-coloured figs than dull or dark ones. This indicates that non-morphological traits (e.g., odour, nutrition, accessibility) may play a stronger role in fig selection, and that large figs can be consumed piecemeal (Lomáscolo et al., 2008).

Carnivores consume fruits from shrubs more frequently than any other mammal group, whereas primates and carnivores consume a similar proportion of climbers to each other. Carnivores such as bears, martens, badgers, and canids are known to often consume fruits from shrubs (Koike & Masaki, 2019). Most primates and some carnivores are arboreal and can climb trees, and since many fruits of climbers are found in the canopy and middle strata, both groups consume them (Nakashima & Do Linh San, 2022). Carnivores also differ from other groups in their consumption of fleshy fruits. Previous studies have similarly reported that dry fruits are rarely detected in their scats (Koike & Masaki, 2019; Mudappa et al., 2010; Nakashima & Do Linh San, 2022). Some primates (Lim et al., 2021) and herbivores also consume dry fruits from the Fabaceae family, which is reflected in their higher dry fruit consumption than carnivores.

Herbivores tend to consume longer and somewhat larger non-fig fruits compared to carnivores and primates. In contrast, carnivores generally consume smaller fruits than both primates and herbivores, although not statistically significant. Herbivores include many mammals that are large (>100 kg body mass), and some bear species are also within this size category (McConkey et al., 2018). However, while herbivores tend to consume larger fruits, bears often handle and consume smaller ones (Koike & Masaki, 2019). This suggests that larger body size in frugivorous mammals does not necessarily correspond to the consumption of larger fruits, which may instead be also influenced by differences in fruit handling ability.

The traditional view that mammals predominantly disperse dull-colored and odorous fruits may also be an oversimplification (Valenta & Nevo, 2020). For instance, carnivores in our dataset, which functioned primarily as seed dispersers, frequently consumed fruits with traits typically associated with avian dispersal, such as brightly and darkly colored fleshy fruits from shrubs and climbers. In contrast, herbivores consumed larger, duller, and more drier fruits predominantly from trees. Primates occupied an intermediate position between these two extremes across nearly all fruit traits, including color, size, fleshiness, and growth form. This intermediate positioning may reflect the remarkable ecological flexibility of primates, facilitated by opposable thumbs and versatile foraging strategies across mammalian lineages.

### 4.3 | Body size and handling ability explain fruit size consumption patterns

The positive relationship between body size and non-fig fruit size aligns with previous studies across mammals (Chen & Moles, 2015; Sivault et al., 2023) and birds (Chen & Moles, 2015; McFadden et al., 2022). However, this relationship was relatively weak, with explanatory power across all positive body-size associations remaining around (R^2^) 0.1. Several factors may explain this weak pattern. First, mammals often have relatively large jaws for their body size, enabling smaller species to consume relatively large fruits (Sivault et al., 2023). Second, our dataset includes both dry and fleshy fruits, which exhibit contrasting relationships with frugivore body size (Chen & Moles, 2015). Finally, the inclusion of both obligate and facultative frugivores, many of which consume fruits opportunistically or seasonally, may obscure size-based patterns (Sridhara et al., 2016). Consistent with this, positive relationships between fruit and mammal body size are strongest when frugivory constitutes more than 50% of the diet, a condition not met by many species in our dataset (Sivault et al., 2023).

In addition to body size, fruit handling ability was important in determining the maximum size of non-fig fruits consumed. We observed a positive relationship between mammal body size with and maximum non-fig fruit diameter among species that handle fruits exclusively with their mouthparts and the mammals handled by forelimbs (Table S10). These mammals, mainly terrestrial herbivores and carnivores, are limited to handling fruits with their mouthparts and with some use of their forelimbs, and therefore they consume fruits only up to a manageable size. For mammals that use forelimbs to handle fruit, there is no positive relationship for mean non-fig fruit size. This indicates that even partial handling capability enables animals to consume larger fruits relative to their body size. A similar pattern occurs in primates, the group with opposable thumbs: because dexterous handling allows them to process fruits of almost any size, for example, smaller primates like lorises consuming bigger size fruits (Fig S11 & S12), fruit size is effectively uncoupled from body size for primates..

The relatively weak relationships we observed in mammals suggest that fruit size may not be strongly constrained by frugivore body size alone. This may reflect the considerable diversity in mammalian feeding strategies and fruit handling abilities compared to birds, and suggests that mammals may be better considered as a set of functionally distinct groups rather than as a single entity. Body size is important to predict the fruit size mammals consume but their diverse handling strategy is also important for determining the maximum size non-fig fruit that mammals consume. Mammals that can use the help of forelimbs or opposable thumbs to manipulate fruits can manipulate fruits of broader range size than those that rely mainly on their mouthparts.

We also found that the relationship between fruit traits and mammalian consumption differed markedly between figs and non-fig fruits. For figs, neither body size nor fruit handling capability imposes strict constraints on fig consumption. Given that figs are soft, not very variable morphologically, and physically easy to process, they can be consumed by a broad range of frugivores regardless of body size. Figs play a distinct ecological and nutritional role providing essential nutrients and often produce massive crop loads (Shanahan et al., 2001), which makes figs very important fruit resources, particularly in tropical regions, where figs are exceptionally abundant in some habitats (Shanahan et al., 2001).

Taken together, these findings highlight the need for a more nuanced perspective on mammalian frugivory rather than approaches that focus on simplistic generalizations. Given the extensive morphological and behavioral diversity across mammalian groups, accounting for functional distinctions like handling ability is essential to understanding the complexity of mammal-frugivore interactions.

## 5 | Caveats and future directions

Ours is the first study to comprehensively examine plant–mammal frugivory interactions at a continental scale across Asia. However, we acknowledge the taxonomic and geographic shortfalls and biases in our dataset, a common feature of species interaction information (Hortal et al., 2015) given the difficulty in recording such information. We also acknowledge that primates are better studied with geographically larger ranges than the other two mammal groups. Furthermore, we focused primarily on morphological traits and could not incorporate other important fruit characteristics, such as nutritional composition or secondary metabolites, quantitative visual traits or volatile compounds (Valenta & Nevo, 2020) due to limited data for these traits for plant species. These traits are known to play a significant role in influencing frugivore fruit selection and should be included in future studies to develop further insights and understanding of mammal–fruit interactions.

## 6 | Conclusion

In conclusion, our synthesis indicates that fruit consumption patterns among Asian mammals are shaped by taxonomic associations and some fruit morphological traits like fruit colour, fruit type and growth habit. Primates and carnivores consumed a greater diversity of fruit genera and shared more genera with each other than with herbivores. Herbivores tended to consume larger fruits and, together with primates, showed preferences for dull-coloured fruits, whereas carnivores consumed more fleshy fruits and fruits from lianas and shrubs across a broader colour spectrum. Among mammals, body size is important in predicting the non-fig fruit size that mammals consume, however fruit handling ability with body size is important in predicting maximum size of non-fig fruits consumed. We also found that mammals tend to follow different patterns of frugivory for figs and non-fig fruits.

These findings demonstrate that comparative analyses of mammalian fruit-diet patterns reveal important functional variation in fruit–mammal interactions and underscore the need for increased field-based research on mammalian frugivory. Our findings also suggest that there may be complementarity among mammalian groups in their functional roles. Frugivorous mammals play central functional roles in mediating both mutualistic and antagonistic interactions (McConkey et al., 2026). Given their central functional roles in mediating these interactions, understanding patterns of fruit consumption and interactions across mammalian groups is particularly important. Across Asia, many primates and large herbivores are threatened by defaunation and other anthropogenic pressures (Corlett, 2007). Therefore, an improved understanding of frugivory patterns is critical for anticipating cascading effects on plant diversity and regeneration dynamics (Gardner et al., 2019).

## Supporting information

Supplementary file contains 12 colour figures and 10 tables

Electronic supplementary file 3

Electronic supplementary file 4

Electronic supplementary file 1

Electronic supplementary file 2

