## Supplementary file contains 12 colour figures and 10 tables for "Body size and fruit handling ability explain frugivory patterns among Asian mammals"

**Supplementary material**

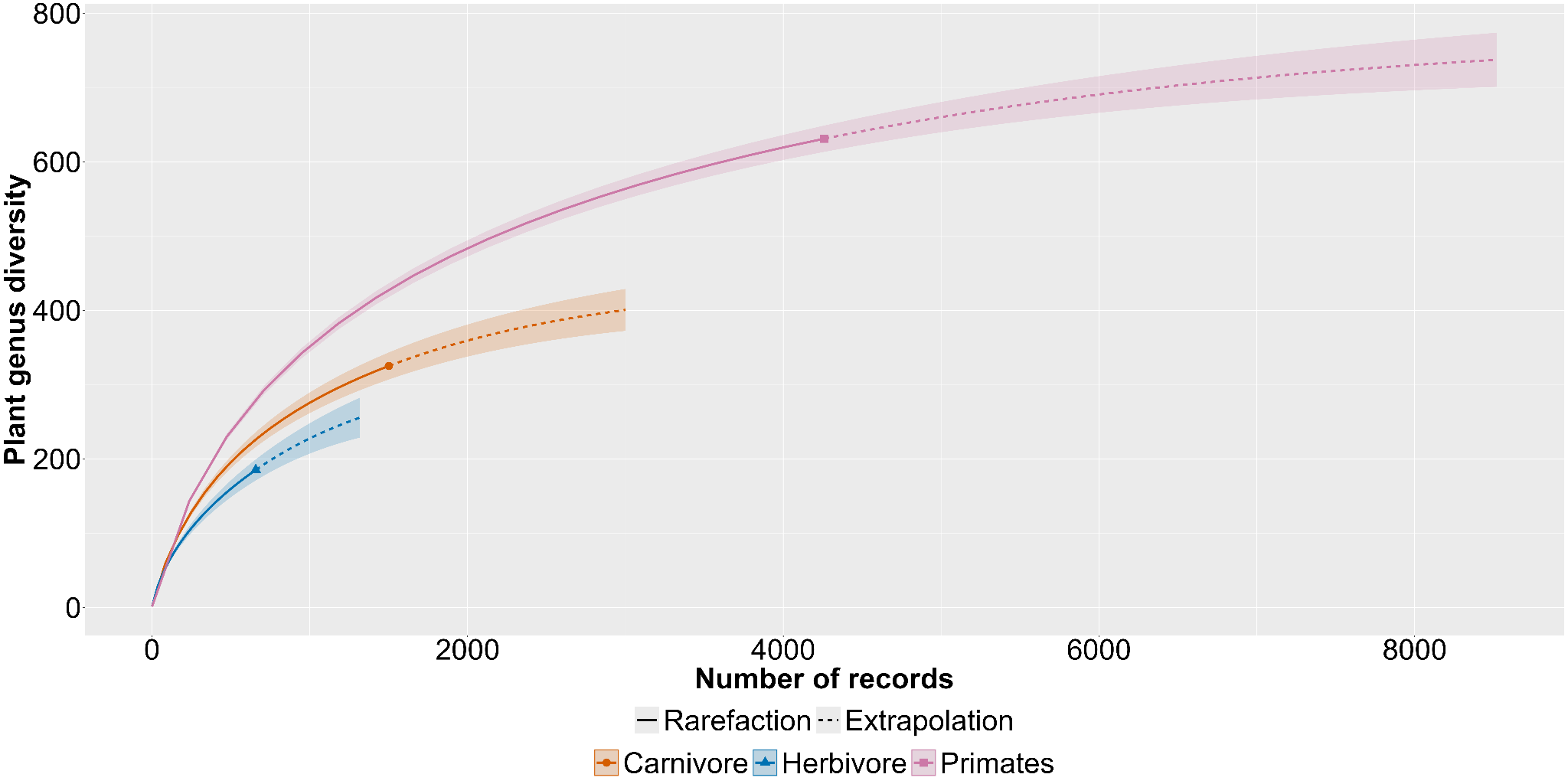

**Figure S1.** Species accumulation curves for plant genera consumed by carnivores, herbivores and primates in Asia, based on the number of independent records (plant genera = 721, number of interaction records = 6419). Coverage was 85.3% for herbivores, 91.9% for carnivores, and 95.6% for primates.

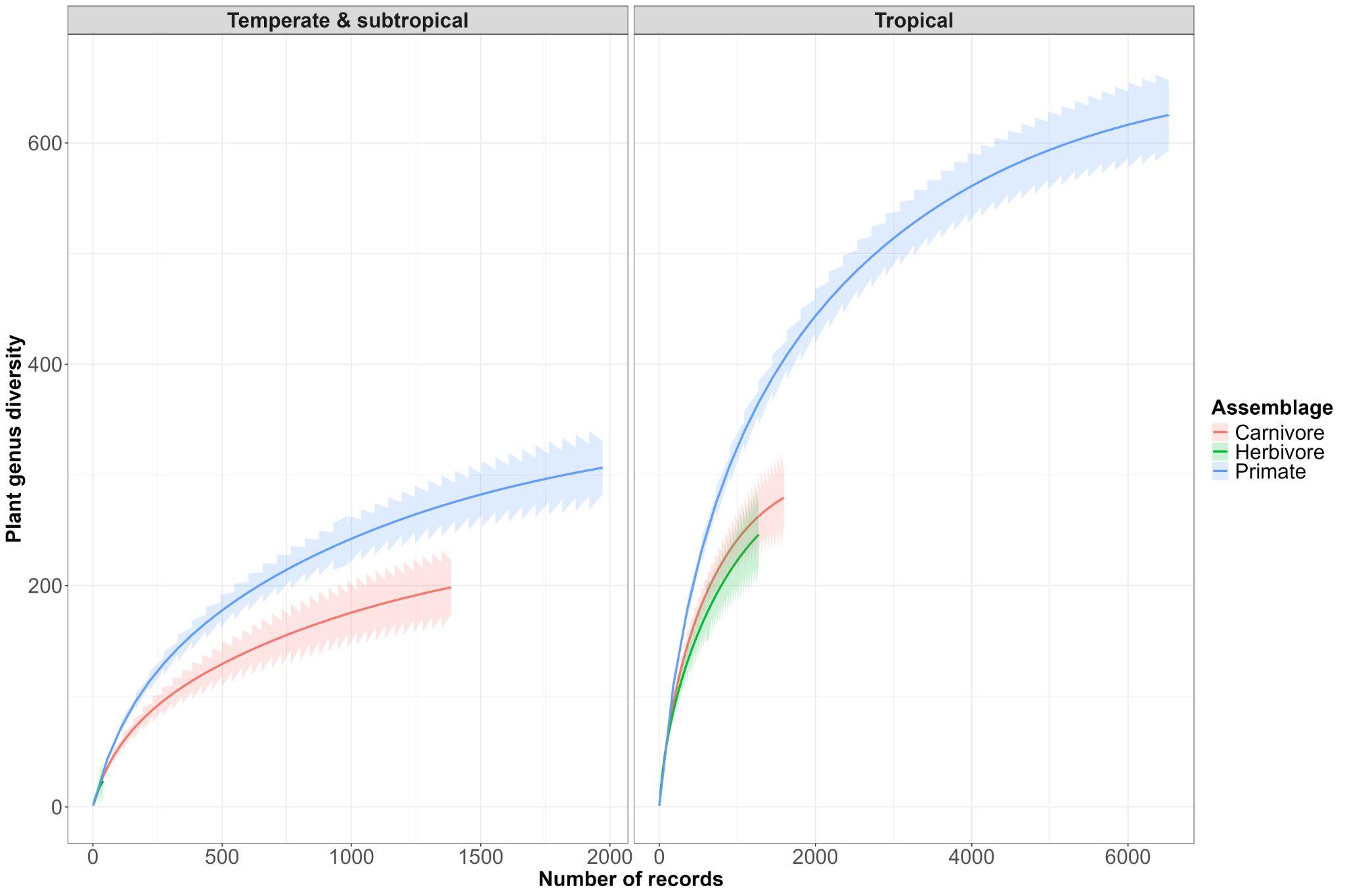

**Figure S2.** Species accumulation curves for plant genera consumed by carnivores, herbivores and primates across different vegetation types in Asia, based on unique records (Temperate & Subtropical: n = 406, plant genera = 296; Tropical: n = 927, plant genera = 587). Coverage value: for Temperate and Subtropical: Carnivore = 90%, Primate = 89%, Herbivore: 45%; Tropical: Primate: 94%, Carnivore: 87 % , Herbivore: 85%).

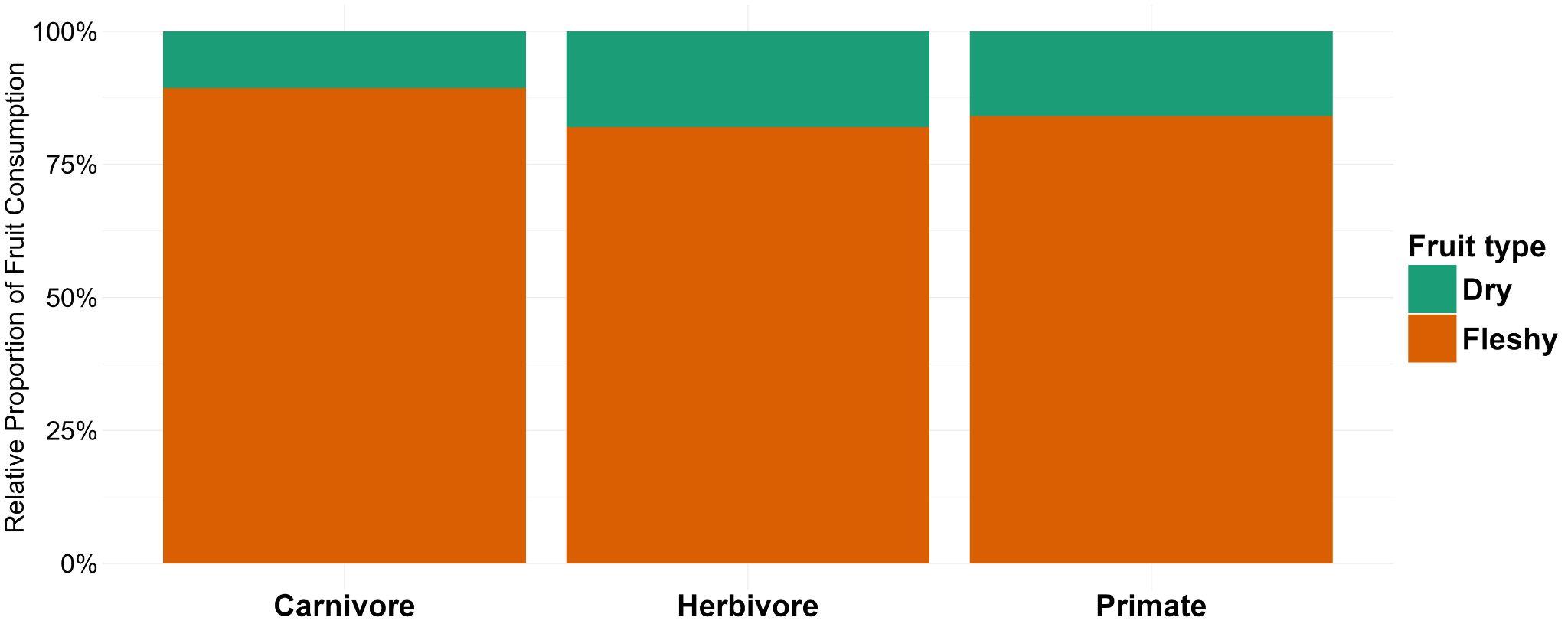

**Figure S3**. Patterns of fruit types consumed by different mammal groups across Asia (Carnivore: 1282, Herbivore: 712, Primate: 3492).

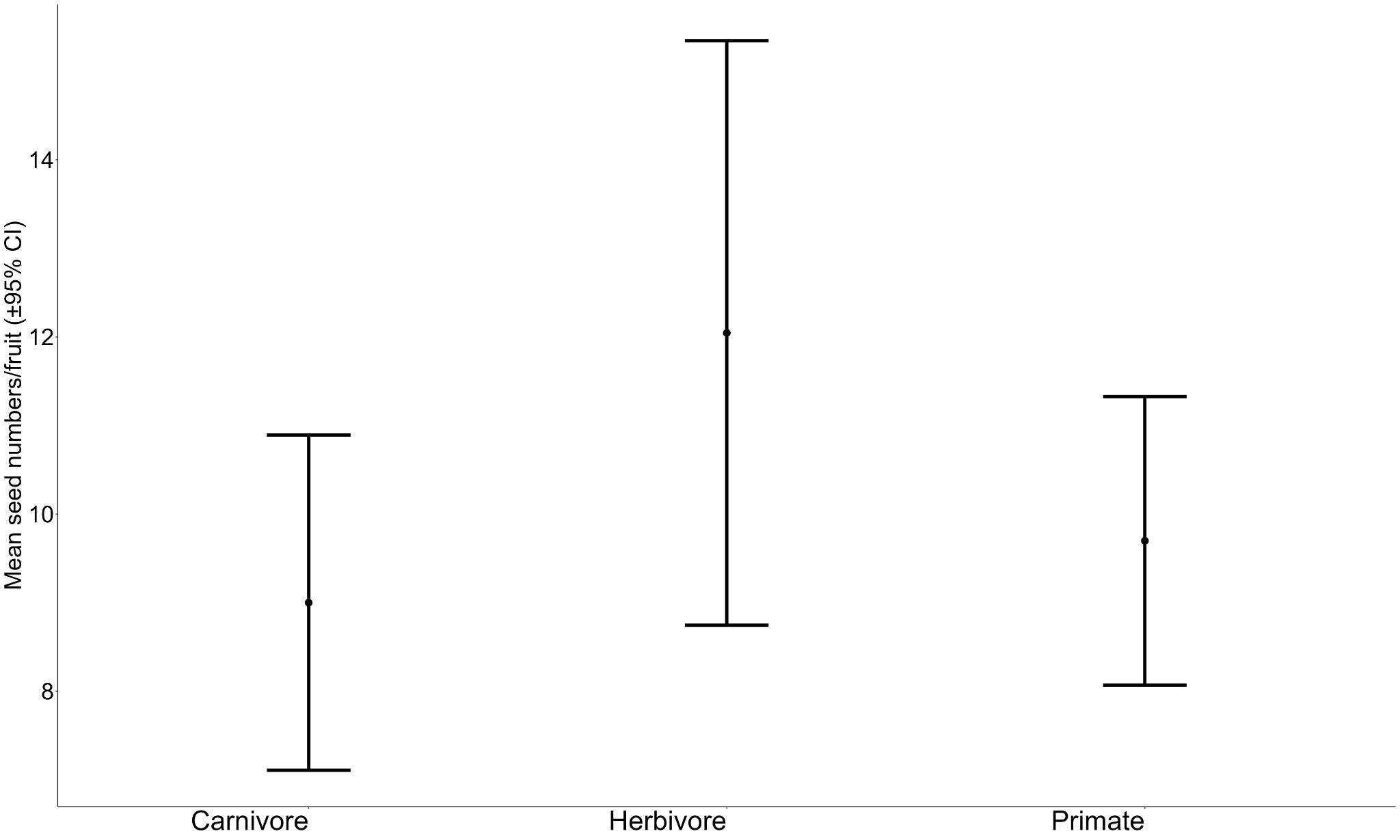

**Figure S4.** Number of seeds per fruit consumed by different mammals across Asia (Carnivore: 1100, Herbivore: 642, Primate: 2983

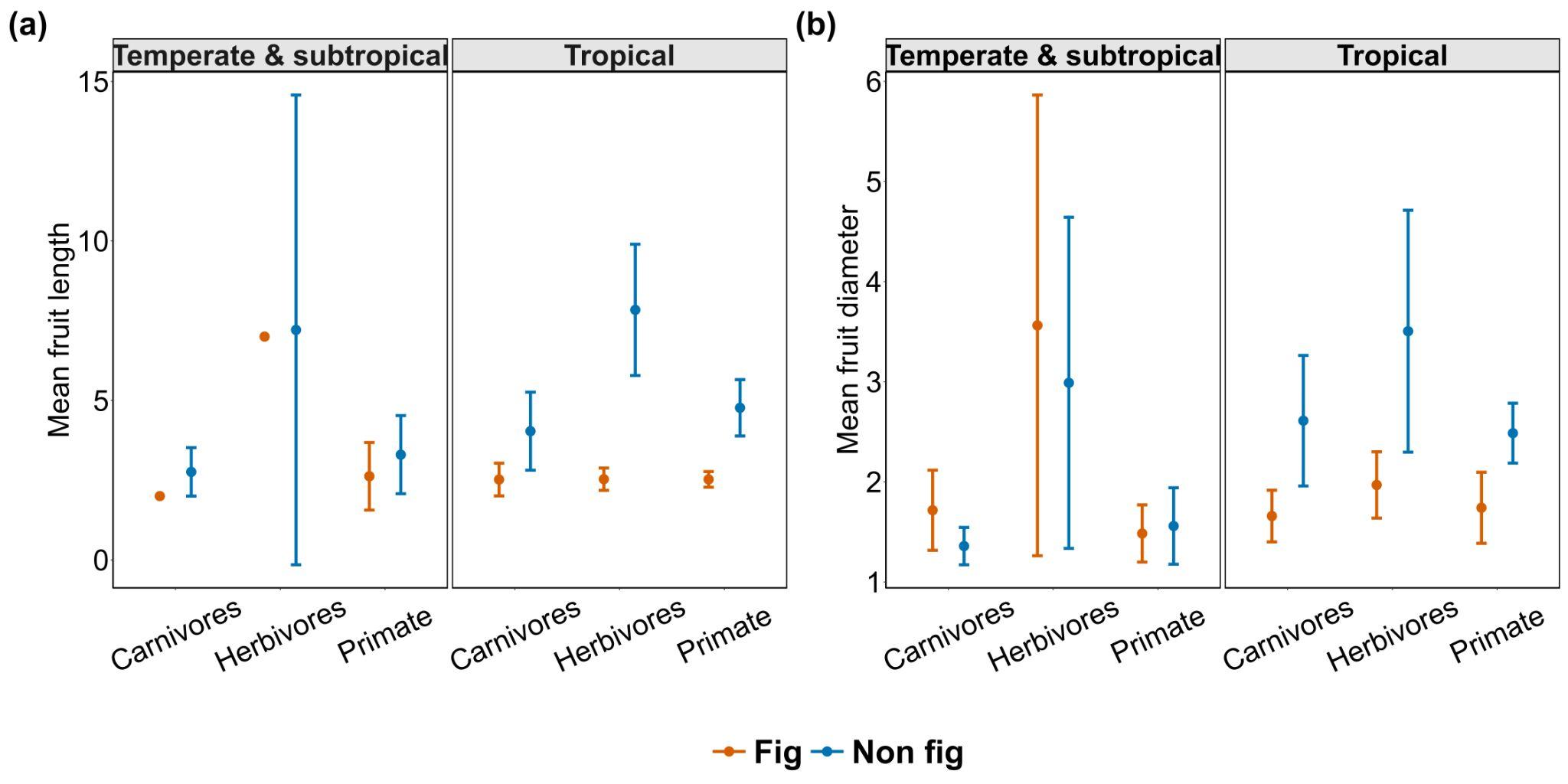

**Fig S5.** Differences in mean fruit length and mean fruit diameter consumed by three mammal groups across vegetation types (a & b), for fig and non-fig fruits. ; for a, fruit length for fig (Temperate & Subtropical: Carnivore = 1, Herbivore = 1, Primate = 4; Tropical: Carnivore = 29, Herbivore = 16, Primate = 71), for non-fig (Temperate & Subtropical: Carnivore = 212, Herbivore = 13, Primate = 326 ; Tropical: Carnivore = 490, Herbivore = 525, Primate = 1911); For b, fruit diameter of fig (Temperate & Subtropical: Carnivore = 8, Herbivore = 4, Primate = 45; Tropical: Carnivore =106, Herbivore = 56, Primate = 419 ), for non-fig (Temperate & Subtropical: Carnivore = 603, Herbivore = 18, Primate = 665; Tropical: Carnivore = 719, Herbivore = 695, Primate = 2816). Error bars are 95% confidence intervals.

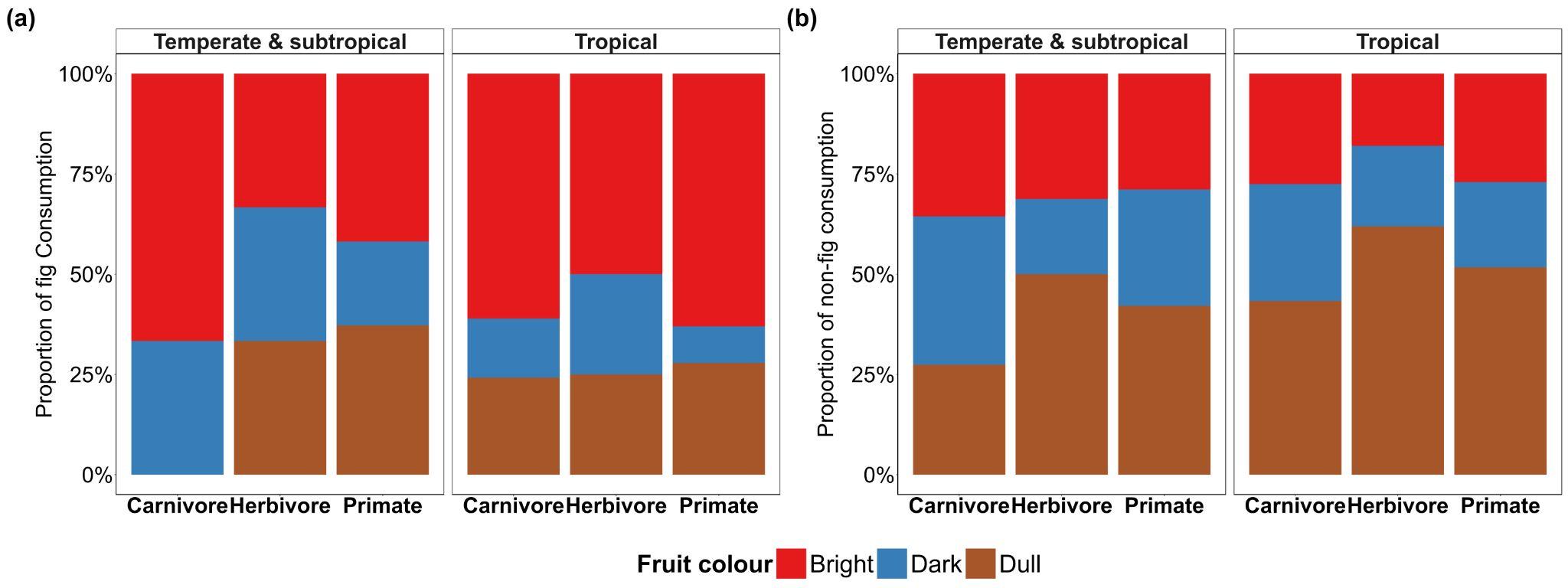

**Fig S6.** Fruit colours of fig and non-fig fruits consumed by different mammal groups across Asia. a) Fig colour (Temperate & Subtropical: Carnivore=8, Herbivore=4, Primates=45, Tropical: Carnivore=106, Herbivore=56, Primate=419), b) non-fig colour (Temperate & Subtropical: Carnivore= 603, Herbivore= 18, Primates= 665; Tropical: Carnivores=719, Herbivore= 695, Primate= 2816)

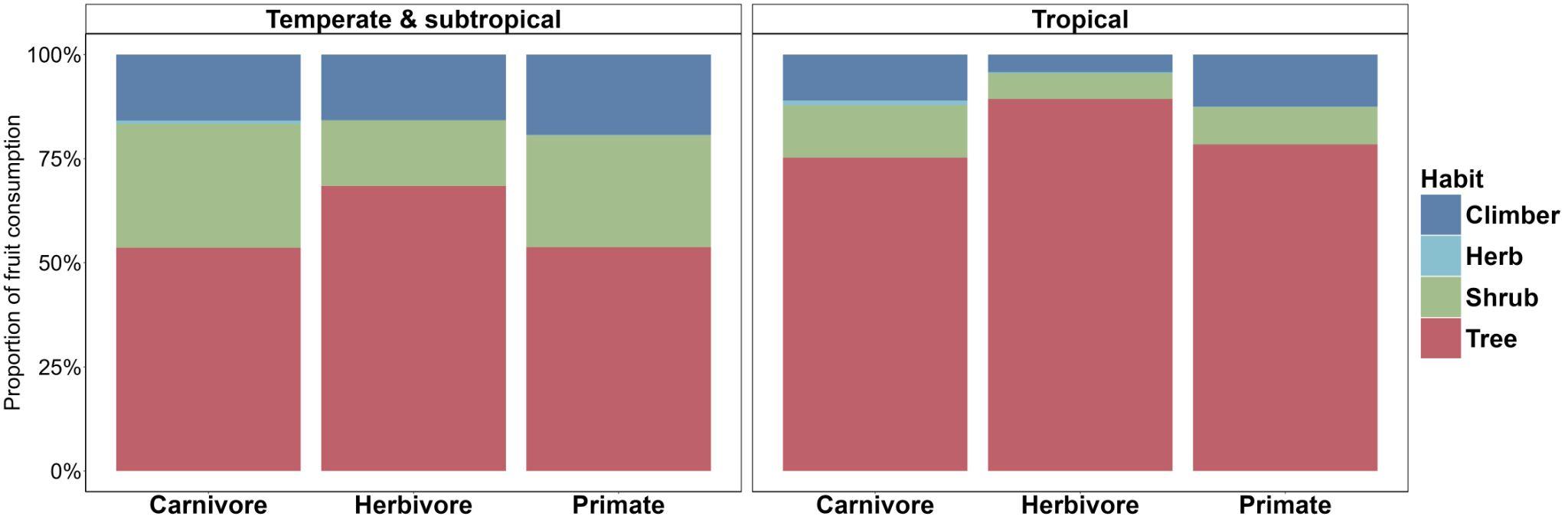

**Fig S7.** Growth habits of the fruits consumed by different mammal groups across different vegetation types. (Temperate & Subtropical: Carnivore = 498, Herbivore = 19, Primate = 695; Tropical: Carnivore = 706, Herbivore = 683, Primate = 2809).

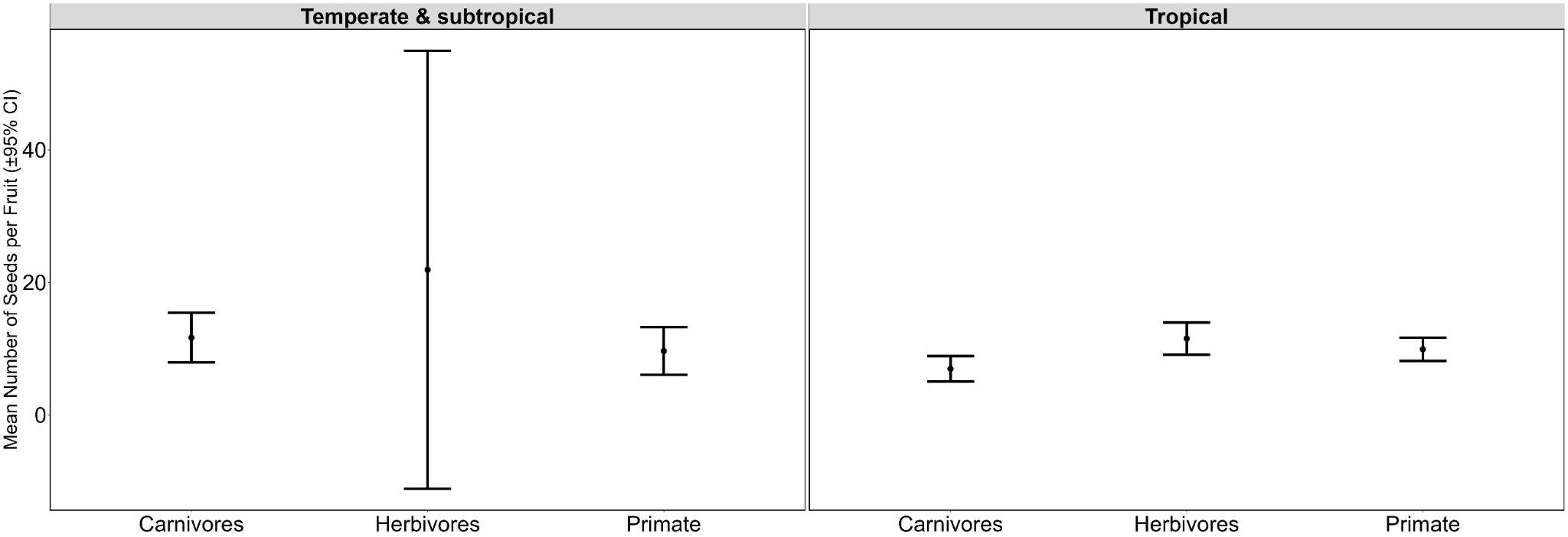

**Figure S8:** Number of seeds per fruit consumed by different mammals across different vegetation types in Asia - Temperate & Subtropical: Carnivore = 491, Herbivore = 16, Primate = 610; Tropical: Carnivore = 609, Herbivore = 629, Primate = 2378.

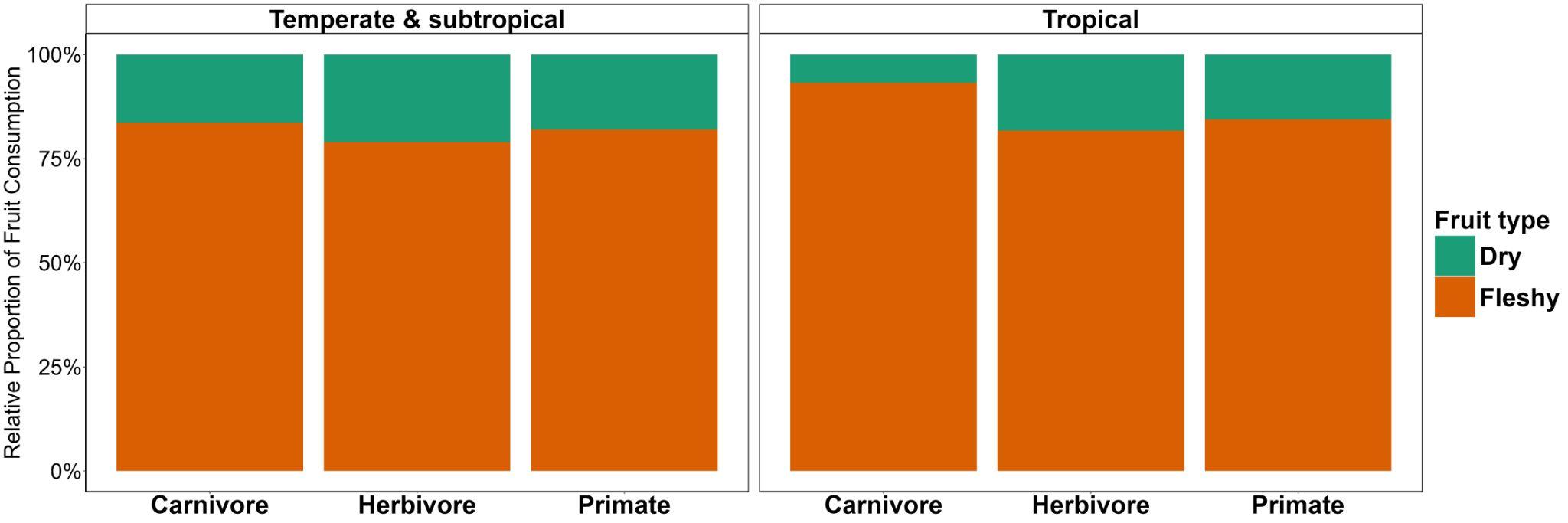

**Figure S9.** Patterns of fruit types consumed by different mammal groups in different vegetation types across Asia - Temperate & Subtropical: Carnivore = 498, Herbivore = 19, Primate = 659; Tropical: Carnivore = 706, Herbivore = 683, Primate = 2809)

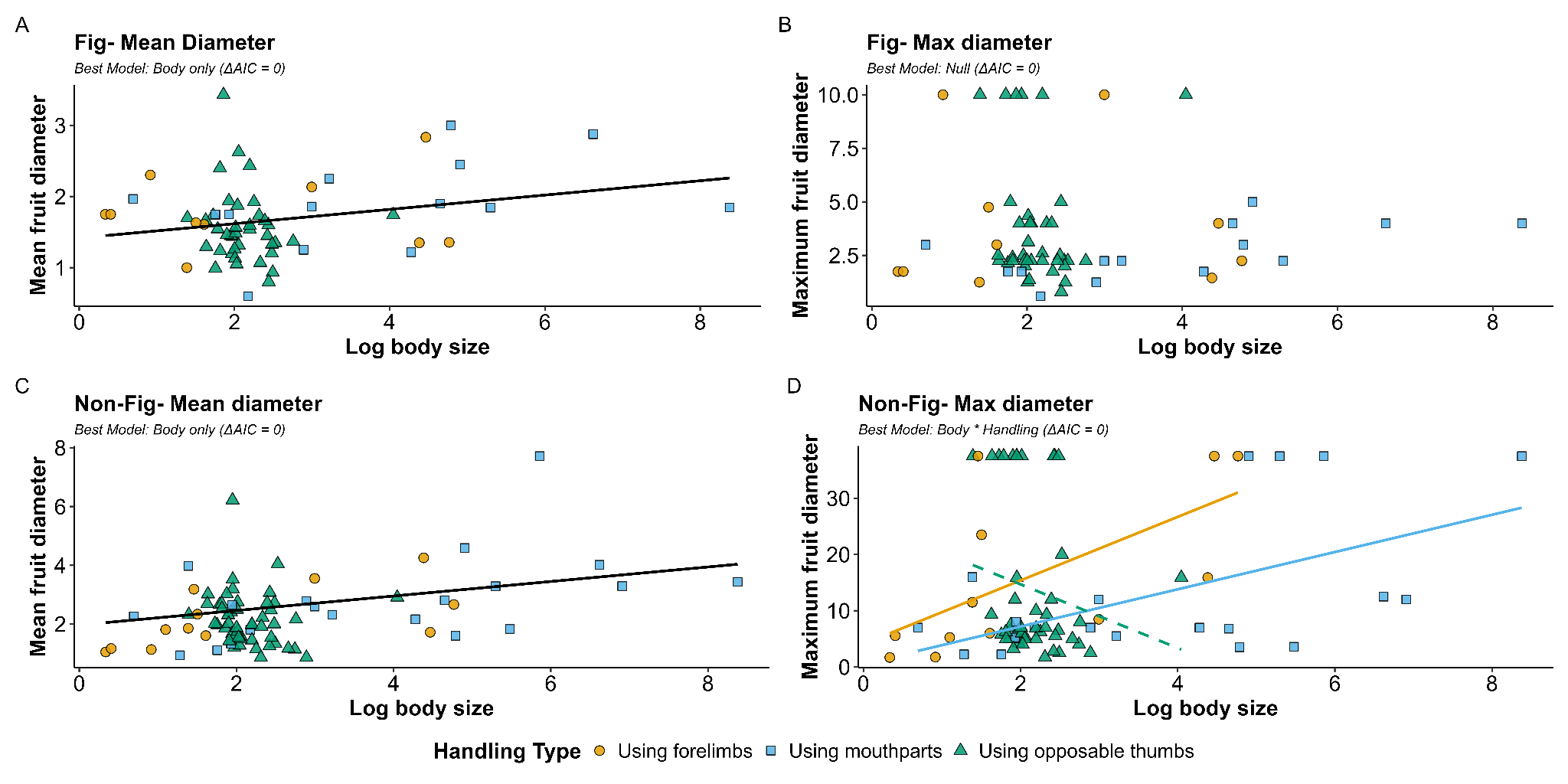

**Fig S10.** Relationships between mammal body size, fruit handling strategy, and consumed fruit diameter. Panels display mean **(A)** and maximum **(B)** fruit diameter for fig species (n= 62), and mean **(C)** and maximum **(D)** diameter for non-fig species (n=78) plotted against log body size (for species with ≥ 10 interaction records). Points represent mammal species categorized by fruit handling ability - forelimbs (orange), mouthparts (blue), opposable thumbs (green). Lines depict best-fit PGLS models selected via AIC, with solid lines indicating significant relationships ( p < 0.05) and dashed lines denoting non-significant trends (p > 0.05).

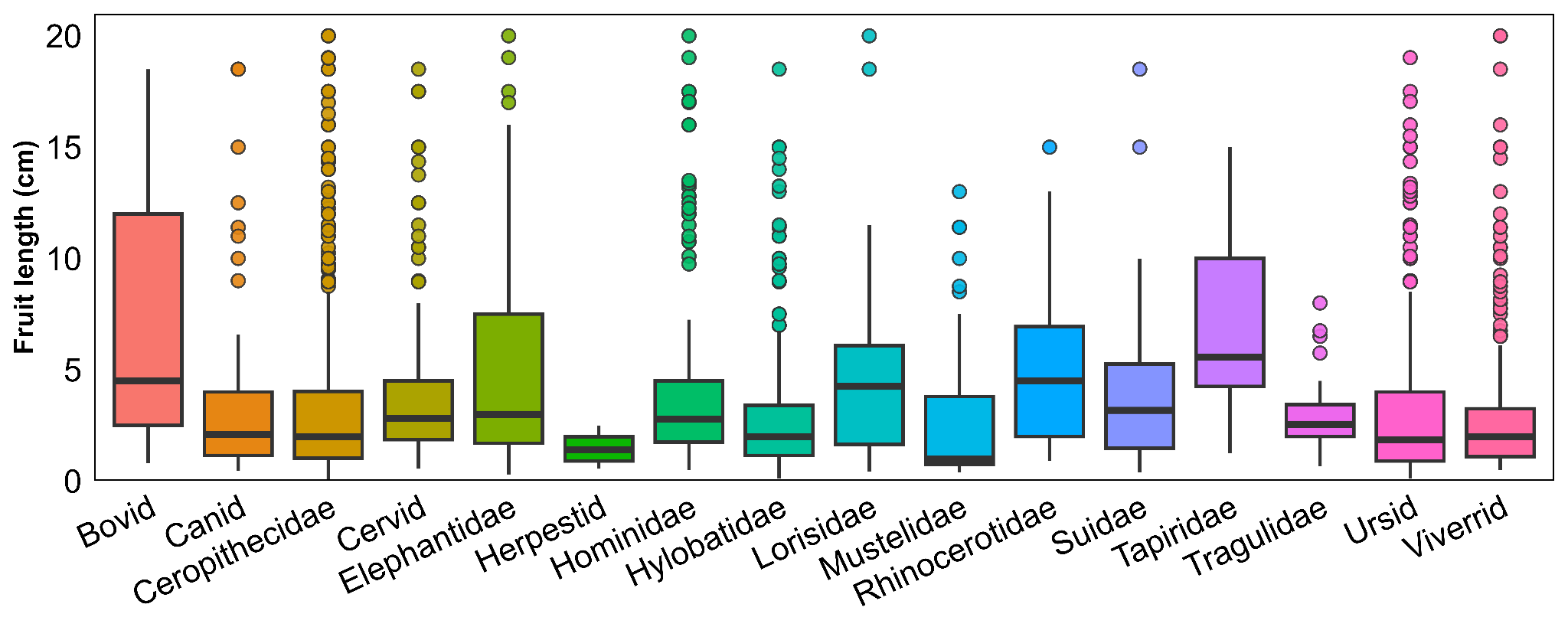

**Figure S11.** Distribution of non-fig fruit length consumed by different mammal families.

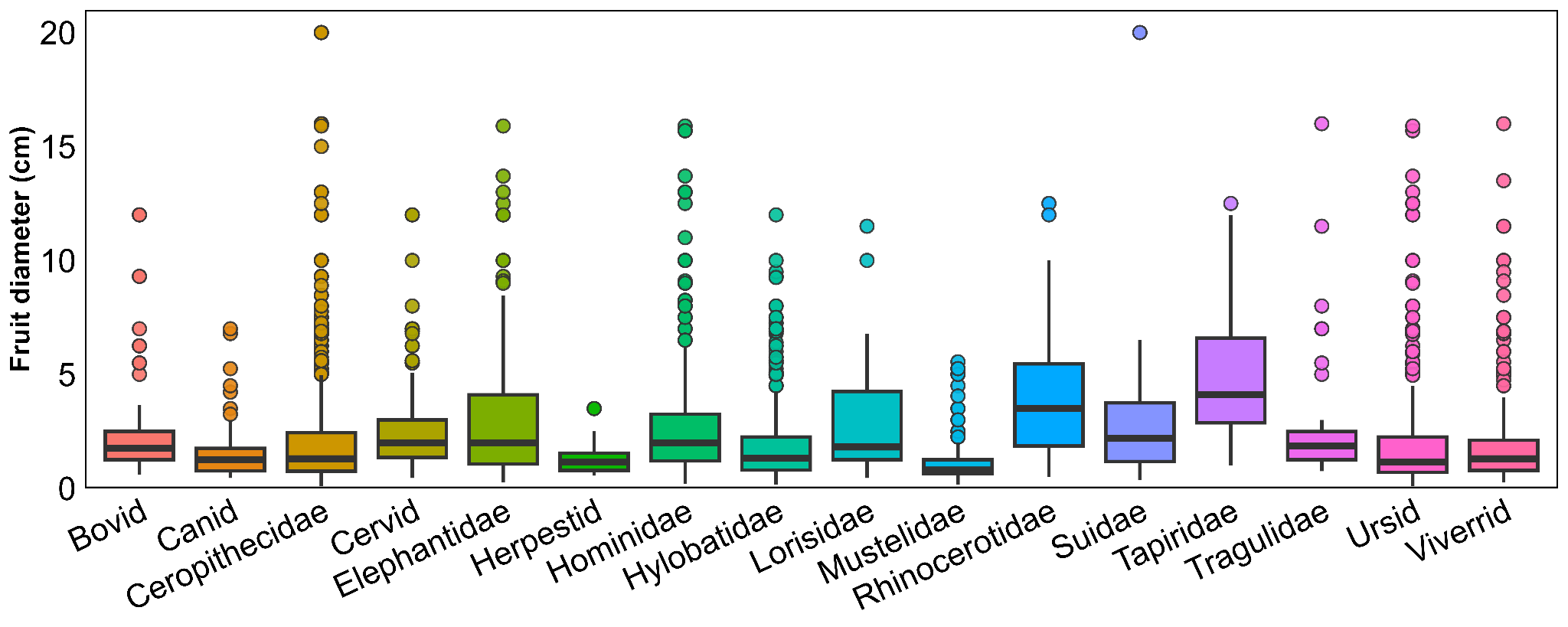

**Figure S12:** Distribution of non-fig fruit diameter consumed by different mammal families.

**Tables**

**Table S1.** Classification of different biomes and ecoregions into three vegetation types <https://ecoregions.appspot.com/>.

| **Vegetation types** | **Biome** |
| --- | --- |
| Temperate & Subtropical | Montane Grasslands & Shrublands |
|  | Temperate Broadleaf & Mixed Forests |
|  | Temperate Conifer Forests |
|  | Tropical & Subtropical Coniferous Forests |
|  | Tropical & Subtropical Moist Broadleaf Forests |
| Tropical | Deserts & Xeric Shrublands |
|  | Flooded Grasslands & Savannas |
|  | Mangroves |
|  | Temperate Broadleaf & Mixed Forests |
|  | Tropical & Subtropical Dry Broadleaf Forests |
|  | Tropical & Subtropical Grasslands, Savannas & Shrublands |

**Table S2. Sample coverage of different fruit taxa consumed by each mammal group for each vegetation type in Asia.**

|  | | Temperate & Subtropical | | Tropical | |
| --- | --- | --- | --- | --- | --- |
|  |  | Taxa consumed | Sample coverage | Taxa consumed | Sample coverage |
| Plant species | Carnivore | 306 | 72% | 428 | 63% |
|  | Primates | 509 | 68% | 1382 | 75% |
|  | Herbivore | 17 | 31% | 339 | 61% |
| Plant genus | Carnivore | 150 | 90% | 220 | 87% |
|  | Primates | 241 | 89% | 528 | 94% |
|  | Herbivore | 15 | 46% | 179 | 85% |
| Plant family | Carnivore | 65 | 97% | 78 | 97% |
|  | Primates | 90 | 98% | 78 | 97% |
|  | Herbivore | 12 | 72% | 61 | 97% |

**Table S3.** R square and F statistics of PERMANOVA tests for group level differences for fruit consumption.

|  | **Sample size** | **R^2^ value** | **Pseudo F value** | ***p*-value** |
| --- | --- | --- | --- | --- |
| **Fig (species)** |  |  |  |  |
| Asia | 114 | 0.045 | 1.8412 | 0.001* |
| Tropical | 108 | 0.044 | 1.64 | 0.007* |
| Temperate & Subtropical | 25 | 0.152 | 1.528 | 0.05 |
| **Non-fig (species)** |  |  |  |  |
| Asia | 2120 | 0.033 | 2.322 | 0.001* |
| Tropical | 1536 | 0.032 | 2.032 | 0.001* |
| Temperate & Subtropical | 697 | 0.08 | 1.779 | 0.001* |
| **Non-fig (Genus)** |  |  |  |  |
| Asia | 721 | 0.0455 | 3.22 | 0.001* |
| Tropical | 585 | 0.042 | 2.66 | 0.001* |
| Temperate & Subtropical | 295 | 0.096 | 2.195 | 0.001* |
| **Non-fig (Family):** |  |  |  |  |
| Asia | 148 | 0.06 | 4.601 | 0.001* |
| Tropical | 123 | 0.045 | 2.78 | 0.001* |
| Temperate & Subtropical | 100 | 0.12 | 2.817 | 0.001* |

**Table S4.** Fruit consumption by carnivores, herbivores, and primates across different vegetation types for tree families, genus and species. The final column presents *p*-values from PERMANOVA tests for mammal group-level differences in fruit consumption. The asterisks indicate significant differences.

|  | **Sample size** | **Uniquely eaten (%)** | | | **Shared (%)** | | | | **PERMANOVA *p*-value** |
| --- | --- | --- | --- | --- | --- | --- | --- | --- | --- |
|  |  | **Herbivore** | **Carnivore** | **Primate** | **Carnivore & Primate** | **Carnivore & Herbivore** | **Herbivore & Primate** | **All Groups** |  |
| Fig (Species) |  | | | | | | | | |
| Tropical | 108 | 5 | 9 | 46 | 20 | 2 | 20 | 13 | 0.014* |
| Temperate | 25 | 0 | 8 | 76 | 4 | 0 | 2 | 4 | 0.064 |
| Non-fig (Species) |  | | | | | | | | |
| Tropical | 1536 | 7 | 8 | 61 | 11 | 1 | 6 | 7 | 0.001* |
| Temperate | 697 | 1 | 29 | 55 | 13 | 0 | 0 | 1 | 0.001* |
| **Non-fig (Genus)** |  | | | | | | | | |
| Tropical | 585 | 3 | 5 | 48 | 15 | 1 | 10 | 17 | 0.001* |
| Temperate & Subtropical | 295 | 1 | 18 | 47 | 29 | 0 | 1 | 3 | 0.001* |
| **Non-fig (Family)** |  | | | | | | | | |
| Tropical | 100 | 6 | 7 | 51 | 12 | 1 | 9 | 15 | 0.001* |
| Temperate & Subtropical | 123 | 1 | 17 | 55 | 21 | 0 | 2 | 3 | 0.001* |

**Table S5.** Details of Kruskal-Wallis One-way ANOVA and the Wilcoxon Matched Pairs Signed Rank tests for fruit size and number of seeds/fruit across Asia and across vegetation types. The asterisk indicates significant differences.

| **Fig/non-Fig** | **Fruit size** | **Pair** | **Scale** | ***p*-value** | **Type of test** |
| --- | --- | --- | --- | --- | --- |
| Fig | Fruit length | NA | Asia | 0.56 | Kruskal-Wallis |
| Non-Fig | Fruit length | NA | Asia | 0.0004* | Kruskal-Wallis |
| Non-Fig | Fruit length | Carnivore & Herbivore | Asia | 0.001* | Wilcoxon Signed Rank Test |
| Non-Fig | Fruit length | Carnivore & Primate | Asia | 0.215 | Wilcoxon Rank Sum Test |
| Non-Fig | Fruit length | Primate & Herbivore | Asia | 0.001* | Wilcoxon Signed Rank Test |
| Non-Fig | Fruit length | NA | Temperate & Subtropical | 0.527 | Kruskal-Wallis |
| Non-Fig | Fruit length | NA | Tropical | 0.005* | Kruskal-Wallis |
| Fig | Fruit length | NA | Temperate & Subtropical | 0.349 | Kruskal-Wallis |
| Fig | Fruit length | NA | Tropical | 0.907 | Kruskal-Wallis |
| Fig | Fruit diameter | NA | Asia | 0.003* | Kruskal-Wallis |
| Fig | Fruit diameter | Herbivore & Primates | Asia | 0.01* | Wilcoxon Signed Rank test |
| Fig | Fruit diameter | Herbivore & Carnivore | Asia | 0.08 | Wilcoxon Signed Rank test |
| Fig | Fruit diameter | Carnivore & Primates | Asia | 0.06 | Wilcoxon Signed Rank test |
| Non-Fig | Fruit diameter | NA | Asia | 0.06 | Kruskal-Wallis |
| Non-Fig | Fruit diameter | NA | Tropical | 0.65 | Kruskal-Wallis |
| Non-Fig | Fruit diameter | NA | Temperate & Subtropical | 0.65 | Kruskal-Wallis |
| Fig | Fruit diameter | NA | Temperate & Subtropical | 0.127 | Kruskal-Wallis |
| Fig | Fruit diameter | NA | Tropical | 0.07 | Kruskal-Wallis |
| Non-fig | Number of seeds/fruit | NA | Asia | 0.530 | Kruskal-Wallis |
| Non-fig | Number of seeds/fruit | NA | Tropical | 0.368 | Kruskal-Wallis |
| Non-fig | Number of seeds/fruit | NA | Temperate & Subtropical | 0.054 | Kruskal-Wallis |

**Table S6.** Summary of fruit colour, fruit type and plant habit of fruits consumed by different mammal groups across Asia and vegetation types, with associated Chi-square test results. The asterisks indicate significant differences.

| **Fruit trait** | **Scale** | **Pair** | **Chi-square value** | **Degrees of freedom** | ***p-*value** |
| --- | --- | --- | --- | --- | --- |
| Fruit colour (Fig) | Asia | Carnivore & Herbivore | 2.76 | 2 | 0.502 |
|  | Asia | Primate & Herbivore | 9.89 | 2 | 0.02* |
|  | Asia | Carnivore & Primate | 2.72 | 2 | 0.502 |
|  | Tropical | NA | 12.55 | 4 | 0.01* |
|  | Temperate & Subtropical | NA | 3.54 | 4 | 0.47 |
| Fruit colour (Non-fig) | Asia | Carnivore & Herbivore | 105.48 | 2 | <0.01* |
|  | Asia | Primate & Herbivore | 32.8 | 2 | <0.01* |
|  | Asia | Carnivore & Primate | 66.36 | 2 | <0.01* |
|  | Tropical | NA | 53.393 | 4 | <0.01* |
|  | Temperate & Subtropical | NA | 27.298 | 4 | <0.01* |
| Fruit type (All) | Asia | NA | 28.637 | 2 | <0.01* |
|  | Tropical | NA | 44.102 | 2 | <0.01* |
|  | Temperate & Subtropical | NA | 0.718 | 2 | 0.698 |
|  | Asia | Carnivore & Herbivore | 20.21 | 1 | <0.01* |
|  | Asia | Primate & Herbivore | 1.73 | 1 | 0.18 |
|  | Asia | Carnivore & Primate | 19.53 | 1 | <0.01* |
| Growth habit (All) | Asia | Carnivore & Herbivore | 116.4 | 2 | <0.01* |
|  | Asia | Primate & Herbivore | 77.64 | 2 | <0.01* |
|  | Asia | Carnivore & Primate | 62.86 | 2 | <0.01* |
|  | Tropical | NA | 83.08 | 6 | <0.01* |
|  | Temperate & Subtropical | NA | 7.41 | 6 | 0.28 |

**Table S7.** Model comparison betweenPGLS and OLS models. Akaike Information Criterion (AIC) values and parameter estimates for Phylogenetic Generalized Least Squares (PGLS) and Ordinary Least Squares (OLS) regression models. For PGLS models, values represent median parameter estimates across 1,000 phylogenetic trees. Analyses were restricted to frugivore species with ≥ 10 interaction records.

| **Dataset** | **Response variable** | **OLS/**  **PGLS** | **Sample size** | **Model** | **AIC** | **Delta AIC** | **R^2^** | **Adjusted R^2^** |
| --- | --- | --- | --- | --- | --- | --- | --- | --- |
| Fig | Mean fruit diameter | OLS | 63 | Log body size only | 103.372 | 0 | 0.063 | 0.048 |
|  | | | | Handling ability only | 105.519 | 2.146 | 0.061 | 0.030 |
|  |  |  |  | Null (y~1) | 105.535 | 2.162 | 0 | 0 |
|  |  |  |  | Log body size x Handling ability | 107.798 | 4.426 | 0.115 | 0.037 |
| Fig | Maximum fruit diameter | OLS | 63 | Null (y~1) | 306.1312 | 0 | 0 | 0 |
|  | | | | Handling ability only | 307.7172 | 1.586 | 0.037 | 0.005 |
|  |  |  |  | Log body size only | 308.1126 | 1.981 | 0.0003 | -0.01 |
|  |  |  |  | Log body size x Handling ability | 312.54 | 6.409 | 0.055 | -0.0275 |
| Non-fig | Mean fruit diameter | OLS | 82 | Log body size only | 235.83 | 0 | 0.168 | 0.157 |
|  | | | | Log body size x Handling ability | 240.37 | 4.542 | 0.202 | 0.149 |
|  |  |  |  | Handling ability only | 246.513 | 10.676 | 0.0752 | 0.051 |
|  |  |  |  | Null (y~1) | 248.921 | 13.084 | 0 | 0 |
| Non-fig | Maximum fruit diameter | OLS | 82 | Log body size x Handling ability | 653.425 | 0 | 0.155 | 0.099 |
|  | | | | Log body size only | 654.274 | 0.848 | 0.0587 | 0.047 |
|  |  |  |  | Null (y~1) | 657.232 | 3.806 | 0 | 0 |
|  |  |  |  | Handling ability only | 660.604 | 7.179 | 0.008 | -0.017 |
| Fig | Mean fruit diameter | PGLS | 62 | Body x Handling | 108.574 | 4.462 | 0.119 | 0.041 |
|  | | | | Body only | 104.111 | 0 | 0.067 | 0.052 |
|  |  |  |  | Handling only | 106.14 | 2.031 | 0.067 | 0.035 |
|  |  |  |  | Null | 106.047 | 1.935 | 0 | 0 |
| Fig | Maximum fruit diameter | PGLS | 62 | Body x Handling | 310.787 | 6.483 | 0.055 | 6.483 |
|  | | | | Body only | 306.287 | 1.984 | 0.0002 | 1.984 |
|  |  |  |  | Handling only | 305.947 | 1.643 | 0.037 | 1.643 |
|  |  |  |  | Null | 304.303 | 0 | 0 | 0 |
| Non-fig | Mean fruit dimeter | PGLS | 78 | Body x Handling | 221.71 | 4.272 | 0.113 | 0.052 |
|  | | | | Body only | 217.439 | 0 | 0.07 | 0.057 |
|  |  |  |  | Handling only | 223.403 | 5.961 | 0.009 | -0.017 |
|  |  |  |  | Null | 220.106 | 2.661 | 0 | 0 |
| Non-fig | Maximum fruit diameter | PGLS | 78 | Body x Handling | 623.542 | 0 | 0.167 | 0.109 |
|  | | | | Body only | 624.474 | 0.932 | 0.066 | 0.054 |
|  |  |  |  | Handling only | 631.462 | 7.919 | 0.004 | -0.022 |
|  |  |  |  | Null | 627.818 | 4.275 | 0 | 0 |

Table S8. Parameter estimates and summary statistics for PGLS models. Model coefficients, *p*-values, R^2^ values, and Pagel’s λ are reported as medians calculated across 1,000 phylogenetic trees. Analyses were restricted to frugivore species with ≥ 10 interaction records.

| **Dataset** | **Response** | **Model**  **specification** | **Term** | **Sample size** | **Coefficients** | ***p* value** | **Median R^2^** | **Adjusted R^2^** | **Median λ** |
| --- | --- | --- | --- | --- | --- | --- | --- | --- | --- |
| Fig | Mean width | Body x Fruit Handling | Using mouthparts | 62 | -0.263 | 0.541 | 0.119 | 0.041 | 1.00E-07 |
| Fig | Mean width | Body x Fruit Handling | Using opposable thumbs | 62 | 0.155 | 0.77 | 0.119 | 0.041 | 1.00E-07 |
| Fig | Mean width | Body x Fruit Handling | Log body size | 62 | 0.034 | 0.741 | 0.119 | 0.041 | 1.00E-07 |
| Fig | Mean width | Body x Fruit Handling | Log body size: Using mouthparts | 62 | 0.085 | 0.501 | 0.119 | 0.041 | 1.00E-07 |
| Fig | Mean width | Body x Fruit Handling | Log body size: Using opposable thumbs | 62 | -0.161 | 0.468 | 0.119 | 0.041 | 1.00E-07 |
| Fig | Mean width | Body only | log body size | 62 | 0.101 | 0.041* | 0.067 | 0.0519 | 1.00E-07 |
| Fig | Mean width | Handling only | Using mouthparts | 62 | 0.125 | 0.576 | 0.067 | 0.0354 | 1.00E-07 |
| Fig | Mean width | Handling only | Using opposable thumbs | 62 | -0.205 | 0.288 | 0.067 | 0.0354 | 1.00E-07 |
| Fig | Max width | Body x Fruit Handling | Using mouthparts | 62 | -3.036 | 0.171 | 0.055 | -0.029 | 1.00E-07 |
| Fig | Max width | Body x Fruit Handling | Using opposable thumbs | 62 | -0.664 | 0.806 | 0.055 | -0.029 | 1.00E-07 |
| Fig | Max width | Body x Fruit Handling | Log body size | 62 | -0.091 | 0.863 | 0.055 | -0.029 | 1.00E-07 |
| Fig | Max width | Body x Fruit Handling | log body size: Using mouthparts | 62 | 0.461 | 0.477 | 0.055 | -0.029 | 1.00E-07 |
| Fig | Max width | Body x Fruit Handling | log body size: Using opposable thumbs | 62 | 0.209 | 0.857 | 0.055 | -0.029 | 1.00E-07 |
| Fig | Max width | Body only | log body size | 62 | -0.031 | 0.901 | 0.0002 | -0.016 | 1.00E-07 |
| Fig | Max width | Handling only | Using mouthparts | 62 | -1.387 | 0.219 | 0.037 | 0.005 | 1.00E-07 |
| Fig | Max width | Handling only | Using opposable thumbs | 62 | -0.203 | 0.832 | 0.037 | 0.005 | 0.651 |
| Non-fig | Mean width | Body x Fruit Handling | Using mouthparts | 78 | 0.360 | 0.672 | 0.113 | 0.052 | 0.651 |
| Non-fig | Mean width | Body x Fruit Handling | Using opposable thumbs | 78 | 0.824 | 0.562 | 0.113 | 0.052 | 0.652 |
| Non-fig | Mean width | Body x Fruit Handling | log body size | 78 | 0.480 | 0.041* | 0.113 | 0.052 | 0.653 |
| Non-fig | Mean width | Body x Fruit Handling | log body size: Using mouthparts | 78 | -0.235 | 0.376 | 0.113 | 0.052 | 0.764 |
| Non-fig | Mean width | Body x Fruit Handling | log body size: Using opposable thumbs | 78 | -0.678 | 0.077 | 0.113 | 0.052 | 0.764 |
| Non-fig | Mean width | Body only | log body size | 78 | 0.247 | 0.019* | 0.069 | 0.057 | 1.00E-07 |
| Non-fig | Mean width | Handling only | Using mouthparts | 78 | 0.05343 | 0.939 | 0.009 | -0.017 | 1.00E-07 |
| Non-fig | Mean width | Handling only | Using opposable thumbs | 78 | -1.041 | 0.462 | 0.009 | -0.017 | 1.00E-07 |
| Non-fig | Max width | Body x Fruit Handling | Using mouthparts | 78 | -3.532 | 0.675 | 0.167 | 0.109 | 1.00E-07 |
| Non-fig | Max width | Body x Fruit Handling | Using opposable thumbs | 78 | 21.931 | 0.054 | 0.167 | 0.109 | 1.00E-07 |
| Non-fig | Max width | Body x Fruit Handling | log body size | 78 | 5.649 | 0.017* | 0.167 | 0.109 | 1.00E-07 |
| Non-fig | Max width | Body x Fruit Handling | log body size: Using mouthparts | 78 | -2.335 | 0.383 | 0.167 | 0.109 | 1.00E-07 |
| Non-fig | Max width | Body x Fruit Handling | log body size: Using opposable thumbs | 78 | -11.3133 | 0.022 | 0.167 | 0.109 | 1.00E-07 |
| Non-fig | Max width | Body only | log body size | 78 | 2.278 | 0.023 | 0.066 | 0.053 | 1.00E-07 |
| Non-fig | Max width | Handling only | Using mouthparts | 78 | -2.627 | 0.588 | 0.004 | -0.021 | 1.00E-07 |
| Non-fig | Max width | Handling only | Using opposable thumbs | 78 | -2.313 | 0.592 | 0.004 | -0.021 | 1.00E-07 |

**Table S9.** Parameter estimates and summary statistics for Ordinary Least Squares (OLS) models. Model coefficients, *p*-values, and R^2^  values are reported for each candidate model. Analyses were restricted to frugivore species with ≥ 10 interaction records.

| **Dataset** | **Response** | **Sample size** | **Model specification** | **Term** | **Estimate** | **Standard error** | ***p* value** | **R^2^** | **Adjusted R^2^** |
| --- | --- | --- | --- | --- | --- | --- | --- | --- | --- |
| Fig | Mean fruit diameter | 62 | Body size only | Log body size | 0.098 | 0.048 | 0.045* | 0.063 | 0.048 |
| Fig | Mean fruit diameter | 62 | Fruit handling | Using mouthparts | 0.125 | 0.222 | 0.575 | 0.061 | 0.03 |
| Fig | Mean fruit diameter | 62 | Fruit handling | Using opposable thumbs | -0.193 | 0.191 | 0.315 | 0.062 | 0.03 |
| Fig | Mean fruit diameter | 62 | Body size x Fruit handling | Log body size | 0.034 | 0.103 | 0.74 | 0.115 | 0.04 |
| Fig | Mean fruit diameter | 62 | Body size x Fruit handling | Using mouthparts | -0.263 | 0.427 | 0.540 | 0.115 | 0.038 |
| Fig | Mean fruit diameter | 62 | Body size x Fruit handling | Using opposable thumbs | 0.204 | 0.524 | 0.698 | 0.115 | 0.038 |
| Fig | Mean fruit diameter | 62 | Body size x Fruit handling | Log body size:Using mouthparts | 0.085 | 0.126 | 0.499 | 0.115 | 0.038 |
| Fig | Mean fruit diameter | 62 | Body size x Fruit handling | Log body size: Using opposable thumbs | -0.183 | 0.225 | 0.418 | 0.115 | 0.038 |
| Fig | Maximum fruit diameter | 62 | Body size only | Log body size | -0.032 | 0.244 | 0.894 | 0.0002 | -0.016 |
| Fig | Maximum fruit diameter | 62 | Fruit handling | Using mouthparts | -1.388 | 1.108 | 0.215 | 0.037 | 0.005 |
| Fig | Maximum fruit diameter | 62 | Fruit handling | Using opposable thumbs | -0.199 | 0.948 | 0.835 | 0.0376 | 0.005 |
| Fig | Maximum fruit diameter | 62 | Body size x Fruit handling | Log body size | -0.091 | 0.524 | 0.862 | 0.055 | -0.027 |
| Fig | Maximum fruit diameter | 62 | Body size x Fruit handling | Using mouthparts | -3.036 | 2.168 | 0.166 | 0.055 | -0.027 |
| Fig | Maximum fruit diameter | 62 | Body size x Fruit handling | Using opposable thumbs | -0.641 | 2.66 | 0.811 | 0.055 | -0.027 |
| Fig | Maximum fruit diameter | 62 | Body size x Fruit handling | Log body size:Using mouthparts | 0.461 | 0.638 | 0.473 | 0.055 | -0.027 |
| Fig | Maximum fruit diameter | 62 | Body size x Fruit handling | Log body size:Using opposable thumbs | 0.201 | 1.146 | 0.861 | 0.055 | -0.027 |
| Non-fig | Mean fruit diameter | 78 | Body size only | Log body size | 0.306 | 0.076 | 0.0001 | 0.168 | 0.157 |
| Non-fig | Mean fruit diameter | 78 | Fruit handling | Using mouthparts | 0.613 | 0.385 | 0.116 | 0.075 | 0.052 |
| Non-fig | Mean fruit diameter | 78 | Fruit handling | Using opposable thumbs | -0.088 | 0.339 | 0.795 | 0.075 | 0.052 |
| Non-fig | Mean fruit diameter | 78 | Body size x Fruit handling | Log body size | 0.303 | 0.187 | 0.109 | 0.202 | 0.15 |
| Non-fig | Mean fruit diameter | 78 | Body size x Fruit handling | Using mouthparts | 0.023 | 0.676 | 0.972 | 0.202 | 0.149 |
| Non-fig | Mean fruit diameter | 78 | Body size x Fruit handling | Using opposable thumbs | 1.111 | 0.889 | 0.215 | 0.202 | 0.149 |
| Non-fig | Mean fruit diameter | 78 | Body size x Fruit handling | Log body Size:Using mouthparts | 0.014 | 0.215 | 0.947 | 0.202 | 0.149 |
| Non-fig | Mean fruit diameter | 78 | Body size x Fruit handling | Log body size:Using opposable thumbs | -0.563 | 0.387 | 0.149 | 0.202 | 0.149 |
| Non-fig | Maximum fruit diameter | 78 | Body size only | Log body size | 2.181 | 0.977 | 0.028* | 0.059 | 0.047 |
| Non-fig | Maximum fruit diameter | 78 | Fruit handling | Using mouthparts | -3.447 | 4.814 | 0.476 | 0.008 | -0.017 |
| Non-fig | Maximum fruit diameter | 78 | Fruit handling | Using opposable thumbs | -3.073 | 4.238 | 0.47 | 0.008 | -0.017 |
| Non-fig | Maximum fruit diameter | 78 | Body size x Fruit handling | Log body size | 5.348 | 2.318 | 0.024* | 0.154 | 0.099 |
| Non-fig | Maximum fruit diameter | 78 | Body size x Fruit handling | Using mouthparts | -4.988 | 8.401 | 0.554 | 0.155 | 0.099 |
| Non-fig | Maximum fruit diameter | 78 | Body size x Fruit handling | Using opposable thumbs | 19.773 | 11.037 | 0.077 | 0.155 | 0.099 |
| Non-fig | Maximum fruit diameter | 78 | Body size x Fruit handling | Log body size:Using mouthparts | -2.035 | 2.668 | 0.448 | 0.155 | 0.099 |
| Non-fig | Maximum fruit diameter | 78 | Body size x Fruit handling | Log body size:Using opposable thumbs | -10.713 | 4.799 | 0.028* | 0.155 | 0.099 |

**Table S10.** Comparison of fruit handling method slopes for Ordinary Least Squares (OLS) and Phylogenetic Generalized Least Squares (PGLS) models. Estimated slopes for each fruit handling method across OLS and PGLS regressions. For PGLS models, values represent median parameter estimates across 1,000 phylogenetic trees. Analyses were restricted to frugivore species with ≥ 10 interaction records.

| **Dataset** | **Response Variable** | **OLS/PGLS** | **Sample size** | **Fruit handling method** | **Estimate** | ***p*-value** |
| --- | --- | --- | --- | --- | --- | --- |
| Fig | Mean diameter | OLS | 10 | Using forelimbs | 0.034 | 0.74 |
| Fig | Mean diameter | OLS | 14 | Using mouthparts | 0.12 | 0.099 |
| Fig | Mean diameter | OLS | 39 | Using opposable thumbs | -0.149 | 0.459 |
| Fig | Maximum diameter | OLS | 10 | Using forelimbs | -0.091 | 0.862 |
| Fig | Maximum diameter | OLS | 14 | Using mouthparts | 0.369 | 0.314 |
| Fig | Maximum diameter | OLS | 39 | Using opposable thumbs | 0.109 | 0.915 |
| Non-fig | Mean diameter | OLS | 12 | Using forelimbs | 0.303 | 0.109 |
| Non-fig | Mean diameter | OLS | 20 | Using mouthparts | 0.317 | 0.004* |
| Non-fig | Mean diameter | OLS | 50 | Using opposable thumbs | -0.26 | 0.445 |
| Non-fig | Maximum diameter | OLS | 12 | Using forelimbs | 5.348 | 0.024* |
| Non-fig | Maximum diameter | OLS | 20 | Using mouthparts | 3.313 | 0.014* |
| Non-fig | Maximum diameter | OLS | 50 | Using opposable thumbs | -5.364 | 0.205 |
| Fig | Mean diameter | PGLS | 10 | Using forelimbs | 0.034 | 0.739 |
| Fig | Mean diameter | PGLS | 14 | Using mouthparts | 0.12 | 0.095 |
| Fig | Mean diameter | PGLS | 38 | Using opposable thumbs | -0.131 | 0.515 |
| Fig | Maximum diameter | PGLS | 10 | Using forelimbs | -0.091 | 0.863 |
| Fig | Maximum diameter | PGLS | 14 | Using mouthparts | 0.369 | 0.314 |
| Fig | Maximum diameter | PGLS | 38 | Using opposable thumbs | 0.118 | 0.909 |
| Non-fig | Mean diameter | PGLS | 12 | Using forelimbs | 0.48 | 0.037* |
| Non-fig | Mean diameter | PGLS | 20 | Using mouthparts | 0.245 | 0.062 |
| Non-fig | Mean diameter | PGLS | 46 | Using opposable thumbs | -0.199 | 0.508 |
| Non-fig | Maximum diameter | PGLS | 12 | Using forelimbs | 5.649 | 0.014* |
| Non-fig | Maximum diameter | PGLS | 20 | Using mouthparts | 3.313 | 0.012* |
| Non-fig | Maximum diameter | PGLS | 46 | Using opposable thumbs | -5.664 | 0.183 |
