## Supplementary material for "Body size and fruit handling ability explain frugivory patterns among Asian mammals": Electronic supplementary file 2

[*Blumea: Biodiversity, Evolution and Biogeography of Plants*](https://repository.naturalis.nl/col/50) *, Volume 22 - Issue 1 p. 15- 20*

170) Elaeocarpus for Flora Malesiana: The E. stipularis Complex, E. nitidus Group & E. barbulatus- MJE COODE, 2001
171) REVISION OF MALESIAN ENDOSPERMUM (EUPHORBIACEAE) WITH NOTES ON PHYLOGENY AND HISTORICAL BIOGEOGRAPHY- Guererro and Welzen 2011
172) <https://portal.cybertaxonomy.org/flora-malesiana/cdm_dataportal/taxon/6318598a-a0ff-4abf-8a41-d7e2526b10b3>
173) Taxonomic revision of the genus Microcos (Malvaceae-Grewioideae) in Peninsular Malaysia and Singapore- Chung and soepadmo 2011
174) <https://portal.cybertaxonomy.org/flora-malesiana/cdm_dataportal/taxon/02e18183-2756-4a96-ab04-269c493d2e26>
175) STUDIES IN ARTOCARPUS AND ALLIED GENERA, V. A REVISION OF PARARTOCARPUS AND HULLETTIA- Jarett 1960
176) THE CIRCUMSCRIPTION, TAXONOMY AND BIOGEOGRAPHY OF PORTERANDIA (RUBIACEAE – GARDENIEAE)- Zahid & Wong 2010
177) Taxonomic notes on Indian Terminalia (Combretaceae)- Chakrabarty et al 2019
178) <https://link.springer.com/chapter/10.1007/978-94-007-1764-0_6>
179) A new variety of Gymnosporia emarginata (Celastraceae) from the Coromandel Coast of Peninsular India- Umamaheshwari et al 2023
180) **Prosopis cineraria as an Unconventional Legumes, Nutrition and Health Benefits:** [**https://www.intechopen.com/chapters/62401**](https://www.intechopen.com/chapters/62401)
181) Dormancy and germination of Coscinium fenestratum (Gaertn.) Colebr. Seeds, Anilkumar et al 2010
182) A REVISION OF PTEROSPERMUM (MALVACEAE: DOMBEYOIDEAE) IN MALESIA: Ganesan et al 2020
183) FLORA MALESIANA PRECURSOR FOR THE TREATMENT OF MORACEAE 8: OTHER GENERA THAN FICUS: CC Berg 2005
184) The genus capparis L in India: Maurya et al 2020
185) Towards a field guide to the trees of the Nee Soon Swamp Forest (VIII): Sapotaceae: Chan et al 2022
186) Comparison of seed dormancy breaking of Eusideroxylon zwageri from

Bali and Kalimantan soaked with sodium nitrophenolate growth

Regulator; Purba et al 2019

187) <https://botany.dnp.go.th/eflora/floraspecies.html?tdcode=07187>
188) **Contrasting seed biology of two ornamental palms: Pygmy Date Palm (Phoenix roebelenii O’Brien) and Fishtail Palm (Caryota urens L.): Prakash et al 2019**
189) Frugivory and seed dispersal of woody species by the Asian elephant (Elephas maximus) in a mid-elevation tropical evergreen forest in India: Jothish 2013
190) The genus Salacia (Celastraceae: Salaciodeae) in peninsular India: Nandikar 2021
191) Mangifera quadrifida (Anacardiaceae), a new record for Singapore: Ganesan 2019
192) <https://www.monaconatureencyclopedia.com/orania-sylvicola-2/?lang=en>
193) Calamus arborescens (Arecaceae): an addition to the flora of India from West Bengal: Mandal & Chowdhury 2021
194) Notes on Musa rubra Kurz (Musaceae) and reduction of M. laterita Cheesman as conspecific: Joe et al 2016
195) TAXONOMIC HISTORY AND IDENTITY OF Musa rubra WALL. EX KURZ., MUSACEAE: Hakkinen 2003
196) Traveset 1995- [**Avoidance by birds of insect-infested fruits of Vaccinium**](https://www.jstor.org/stable/3545962)
197) <https://portal.cybertaxonomy.org/flora-malesiana/cdm_dataportal/taxon/5977f683-ec0b-4bbf-a66e-3a08a9dde226>
198) <https://tropical.theferns.info/viewtropical.php?id=Aglaia+odoratissima>
199) <https://tropical.theferns.info/viewtropical.php?id=Aglaia+speciosa>
200) <https://www.nparks.gov.sg/florafaunaweb/flora/3/5/3509>
201) <https://www.mybis.gov.my/art/277>
202) Systematics and biogeography of Elaeocarpus (Elaeocarpaceae)- Sook-Ngoh PHOON 2015
203) <https://seedsofborneo.com/2018/12/07/mangifera-torquenda-rodent-dispersed/>
204) <https://tropical.theferns.info/viewtropical.php?id=Mangifera+cochinchinensis>
205) <https://www.nparks.gov.sg/florafaunaweb/flora/5/6/5626>
206) [https://web.archive.org/web/20080412002556/http://www.pendernursery.com/Catalog/Detail/callicarpashikokiana.html](https://web.archive.org/web/20080412002556/http:/www.pendernursery.com/Catalog/Detail/callicarpashikokiana.html)
207) <http://www.efloras.org/florataxon.aspx?flora_id=2&taxon_id=200011968>
208) <https://www.philippineplants.org/Families/Meliaceae.html>
209) <https://portal.cybertaxonomy.org/flora-malesiana/cdm_dataportal/taxon/f2f7993e-d347-427e-ab42-e703c5e1e72f>
210) <https://www.asianplant.net/Meliaceae/Aglaia_elliptica.htm>
211) <https://www.asianplant.net/Lamiaceae/Callicarpa_pentandra.htm>
212) <https://apps.worldagroforestry.org/treedb/AFTPDFS/Mangifera_odorata.PDF>
213) Tarszisz, E., Tomlinson, S., Harrison, M.E., Morrogh-Bernard, H.C. and Munn, A.J., 2018. Gardeners of the forest: effects of seed handling and ingestion by orangutans on germination success of peat forest plants. *Biological Journal of the Linnean Society*, *123*(1), pp.125-134.
214) Revision of the Sapotaceae of the Malaysian area in a wider sense. XXIII. Palaquium Blanco by P. van Royen, 1960
215) NEW OR NOTEWORTHY PHILIPPINE PLANTS, By Elmer D. Merrill,
216) StyracaceaeC.G.G.J. van SteenisBuitenzorg) , Flora malesiana
217) A MONOGRAPH OP THE GENUS NEESIA * Blume , REINWARDTIA Published by Herbarium Bogoriense, Kebun Raya Indonesia Volume 5, Part 4, p.p. 481-508 (I960)
218) A Monograph of Ebenaceae vol 12, W.P. Hiern
219) Chong et al.: Tree Flora of Nee Soon (I): Lauraceae
220) Taxonomic review of Polyosma (Escalloniaceae) in Papuasia O.K. Paul1,2,3, B.J. Conn2,3, M.J. Henwood3
221) Importance and Potentiality of Underutilized Lakoocha (Artocarpus lakoocha Roxb) Fruit of Tripura, Islam et al 2018
222) A revision of Cryptocarya (Lauraceae) from Thailand and Indochina R.P.J. de Kok
223) Seed Size Variation of Trees and Lianas in a Tropical Forest of Southeast Asia: Allometry, Phylogeny, and Seed Trait - Plant Functional Trait Relationships, Pothasin et al 2022
224) A taxonomic revision of Garcinia section Xanthochymus (Clusiaceae) in Thailand, Chatchai Ngernsaengsaruay1,2 , Pichet Chanton1 , Minta Chaiprasongsuk1 and Nisa Leksungnoen
225) A taxonomic revision of Garcinia section Discostigma (Clusiaceae) in Thailand Chatchai Ngernsaengsaruay1,2 , Pichet Chanton3 , Weereesa Boonthasak4 , Wanwisa Bhuchaisri5
226) ILEX IN TAIWAN AND THE LIUKIU ISLANDS SlIIL-YING Hu
227) A new combination and new records of Tetrastigma (Vitaceae) from ThailandPHONGSAKORN KOCHAIPHAT1, ANNA TRIAS-BLASI2 & PIMWADEE PORNPONGRUNGRUENG1,*
228) A revision of Syzygium Gaertn. (Myrtaceae) in Indochina (Cambodia, Laos and Vietnam)
229) The Menispermaceae of Malesia and Adjacent Areas: VI* PYCNARRHENA, MACROCOCCULUS & HAEMATOCARPUS L. L. FORMAN
230) STUDIES IN MALESIAN VITACEAE VII. The genus Tetrastigma in the Malay Peninsula A. LATlFF
